# Ferroptosis Integrates Mitochondrial Derangements and Pathological Inflammation to Promote Pulmonary Hypertension

**DOI:** 10.1101/2023.01.19.524721

**Authors:** Felipe Kazmirczak, Neal T. Vogel, Sasha Z. Prisco, Michael T. Patterson, Jeffrey Annis, Ryan T. Moon, Lynn M Hartweck, Jenna B. Mendelson, Minwoo Kim, Natalia Calixto Mancipe, Todd Markowski, LeAnn Higgins, Candace Guerrero, Ben Kremer, Madelyn L. Blake, Christopher J. Rhodes, Jesse W. Williams, Evan L. Brittain, Kurt W. Prins

**Affiliations:** Minneapolis Heart Institute, Minneapolis, MN; Lillehei Heart Institute, Cardiovascular Division, Department of Medicine, University of Minnesota, Minneapolis, MN; Center for Immunology, University of Minnesota, Minneapolis, Minnesota; Division of Cardiovascular Medicine, Department of Medicine, Vanderbilt University Medical Center, Nashville, TN; Minnesota Supercomputing Institute, University of Minnesota, Minneapolis, MN; Center for Metabolomics and Proteomics, Department of Biochemistry, Molecular Biology, and Biophysics, University of Minnesota; National Heart and Lung Institute, Imperial College, London, UK; Department of Integrative Biology and Physiology, University of Minnesota, Minneapolis, Minnesota

## Abstract

**Background:** Mitochondrial dysfunction, characterized by impaired lipid metabolism and heightened reactive oxygen species (ROS) generation, results in lipid peroxidation and ferroptosis. Ferroptosis is an inflammatory mode of cell death that promotes complement activation and macrophage recruitment. In pulmonary arterial hypertension (PAH), pulmonary arterial endothelial cells (PAEC) exhibit cellular phenotypes that promote ferroptosis. Moreover, there is ectopic complement deposition and inflammatory macrophage accumulation in the pulmonary vasculature. However, the effects of ferroptosis inhibition on these pathogenic mechanisms and the cellular landscape of the pulmonary vasculature are incompletely defined.

**Methods:** Multi-omics and physiological analyses evaluated how ferroptosis inhibition modulated preclinical PAH. The impact of AAV1-mediated expression of the pro-ferroptotic protein ACSL4 on PAH was determined, and a genetic association study in humans further probed the relationship between ferroptosis and pulmonary hypertension (PH).

**Results:** Ferrostatin-1, a small-molecule ferroptosis inhibitor, mitigated PAH severity in monocrotaline rats. RNA-seq and proteomics analyses demonstrated ferroptosis was associated with PAH severity. RNA-seq, proteomics, and confocal microscopy revealed complement activation and pro-inflammatory cytokines/chemokines were suppressed by ferrostatin-1. Additionally, ferrostatin-1 combatted changes in endothelial, smooth muscle, and interstitial macrophage abundance and gene activation patterns as revealed by deconvolution RNA-seq. Ferroptotic PAEC damage associated molecular patterns restructured the transcriptomic signature, mitochondrial morphology, and promoted proliferation of pulmonary artery smooth muscle cells, and created a pro-inflammatory phenotype in monocytes *in vitro*. AAV1-*Acsl4* induced an inflammatory PAH phenotype in rats. Finally, single-nucleotide polymorphisms in six ferroptosis genes identified a potential link between ferroptosis and PH severity in the Vanderbilt BioVU repository.

**Conclusions:** Ferroptosis promotes PAH through metabolic and inflammatory mechanisms in the pulmonary vasculature.

## Introduction

The accumulation of iron-dependent lipid peroxides induces ferroptosis, a nonapoptotic form of cell death that integrates metabolic derangements, oxidative stress, and inflammation^1^. Multiple organelles regulate ferroptosis, but the mitochondria may be the most important because mitochondria both initiate and propagate ferroptosis^1^. Altered ferroptosis homeostasis occurs in multiple disease states including cancer, sepsis, cardiovascular diseases, and aging, showing the broad implications this mode of cell death has in medicine^1^. The discovery of ferrostatin-1^2^ and liproxstatin-1^3^, two small molecule inhibitors of ferroptosis, greatly expanded our knowledge of ferroptosis as these molecules exert therapeutic effects in a wide-array of conditions^1^. In the systemic vasculature, ferroptosis induces endothelial cell dysfunction and accelerates atherosclerosis development in mice, and importantly this is suppressed by ferrostatin-1 treatment^4^. In pulmonary arterial hypertension (PAH), pulmonary arterial endothelial cell (PAEC) dysfunction, characterized by disrupted mitochondrial function and impaired iron and lipid metabolism^5^, sets up a pro-ferroptotic environment. This hypothesis is supported by the observation that small molecule and genetic interventions that suppress ferroptosis reduce PAH severity in rodents^6–8^. However, the timing and therapeutic potential of antagonizing ferroptosis in PAH requires further investigation. Certainly, previous rodent studies show ferroptosis suppression reduces PAH severity, but these studies intervene with ferrostatin-1 immediately before monocrotaline injection^6^ or lenti-virus mediated overexpression of the anti-ferroptotic protein peroxiredoxin-6 72 hours prior to monocrotaline injection^7^. At present, the impact of ferroptosis inhibition on pathological pulmonary vascular remodeling after PAH has developed are unknown, and this has important translational implications for treating PAH with ferroptosis suppressing approaches.

In addition to being driven by disrupted metabolism, ferroptosis could promote adverse pulmonary vascular remodeling via modulation of the innate immune system and signaling to surrounding cells in the pulmonary vasculature. Ferroptotic cells release damage associated molecular patterns (DAMPs)^9^, which promote ectopic complement deposition^10^, a finding present in rodent and human PAH^11^. DAMPs also recruit monocyte derived macrophages to areas of tissue damage^12^, which can further amplify the deleterious pro-inflammatory state. In rodent and human PAH, monocyte-derived inflammatory interstitial macrophage numbers are increased in the pulmonary vasculature, and blocking interstitial macrophage invasion into the lungs reduces PAH severity^13^. At present, the direct effects of ferroptotic PAEC DAMPs on macrophage/monocyte biology are unknown. Moreover, the impact of ferroptotic DAMPs on pulmonary artery smooth muscle cells (PASMC), a cell type that regulates pulmonary vascular disease due to heightened proliferation and increased contractility^14^, pathophysiology are unexplored.

To address these important unknowns, we employed mechanistic and translational studies using cell culture-based experiments, physiological and multi-omics evaluations in rodents, and a genetics association study in humans to define the role of ferroptosis in PAH. In particular, we performed a drug intervention rodent study and used quantitative lung proteomics, bulk and deconvolution RNA-sequencing, histological examinations, and lung metabolomics to evaluate how ferroptosis inhibition impacted PAH pathobiology. Next, we determined how AAV1-mediated overexpression of the pro-ferroptotic protein acyl-CoA synthetase long-chain family member 4 (ACSL4)^15^ in PAECs *in vivo* modulated pulmonary vascular disease burden to test the hypothesis that a pro-ferroptotic genetic intervention is sufficient to induce PAH. To gain mechanistic insights and probe potential cell-cell communication via released DAMPs, we defined how ferroptotic PAECs modulated PASMC and monocyte biology *in vitro*. Finally, we performed a human genetics association study to potentially link single nucleotide polymorphisms in ferroptosis genes with PH severity.

## Methods

### Data Availability Statement

Data sets, analysis, and study materials will be made available upon request to other researchers for the purpose of the results or replicating the procedures. All data of OMIC experiments are publicly available. The RNA-sequencing (RNA-seq) data are available at the NCBI Gene Expression Omnibus with the accession numbers GSE277179 and GSE277180. The mass spectrometry (MS) proteomics data are available at Figshare: DOI:10.6084/m9.figshare.27078124

### Rodent Studies

Small-molecule mediated ferroptosis inhibition in the monocrotaline (MCT, Sigma-Aldrich) rat PAH model was evaluated in male Sprague-Dawley (Charles River) rats. Rodents were randomly assigned (coin flip by lead investigator) to one of three groups: 1. Control rats injected with phosphate buffered saline (*n*=5), 2. Rats injected with 60 mg/kg MCT and then treated with daily intraperitoneal injections of vehicle (2% dimethyl sulfoxide, 50% polyethylene glycol, 5% Tween 80, and 43% double distilled water) starting two weeks after MCT injection (*n*=10), and 3. MCT rats treated with daily intraperitoneal ferrostatin-1 (1 mg/kg, Selleck Chemicals) starting two weeks after MCT injection (*n*=10). End-point analysis was performed 24 days post MCT injection with primary physiological read-outs being right ventricular systolic pressure (RVSP) and effective arterial elastance (Ea). AAV1-GFP and AAV1-*Acsl4* at a dose of 0.3×10^11^ vector genomes were delivered via intratrachael injection the same day as low dose MCT (30 mg/kg). The end-point studies for viral study were conducted 24 days post treatment. Females were not evaluated due to the more robust phenotype observed in males. Sample size was estimated at *n*=10 based on previous experience with phenotypic variation and statistical power^16^. For AAV treatments, interim analysis was performed after *n*=5 rodents per group, and statistical differences in RVSP and Ea were observed so the study was terminated. No rodents were excluded. All rodent studies were approved by the University of Minnesota Institutional Animal Care and Use Committee

### Cardiovascular Phenotyping

Echocardiography and closed-chest pressure-volume loop analyses quantified RV function and pulmonary vascular disease as we have previously described^16,17^.

### RNAsequencing

RNA from rat lung tissue was isolated from *n*=4 control, *n*=4 MCT-Vehicle and *n*=4 MCT-Fer-1 using PureLink RNA Mini kit (ThermoFisher) with DNase. RNA sequencing and library preparations were conducted at the University of Minnesota Genomics Center via Illumina NovaSeq 6000 with 20 million reads per sample. PASMC cells were lysed and RNA was isolated using Purelink RNA Mini Kit (Invitrogen) according to kit instructions. RNA was submitted to University of Minnesota Genomics Center for library creation (TruSeq Stranded mRNA, converted to Aviti compatible with Element Adept Rapid PCR + kit) and >250×10^6^ reads were generated.

### Lung Mitochondrial Proteomics Analysis

Mitochondrial enrichments (Abcam) from lung tissue were performed according to manufactures instructions and were subjected to TMT16-plex (ThermoFisher Scientific) labeling and quantitative proteomics using Proteome Discover Software as previously described^16,17^. The lung mitochondrial-enriched proteome in *n*=5 control, *n*=5 MCT-Vehicle, and *n*=6 MCT-Fer-1 animals were analyzed.

### Lung Metabolomics/Lipidomics

Global metabolomics analysis of frozen lung specimens on *n*=5 Control, *n*=5 MCT-Vehicle, and *n*=5 MCT-Fer-1 was performed at the University of Minnesota Center for Metabolomics and Proteomics using the Biocrates’ MxP® Quant 500 kit. Samples were homogenized in a 2.0 mL Precelly standard tubes. Samples were centrifuged at 10,000*g* for 5 minutes at 4°C and supernatant was collected. 10µL of the extract was loaded and classes of metabolites were determined using 50µL of 1:1:1:0.16 water:EtOH:pyridine:phenyl isothiocyanate solution and incubated for an hour. ABSciex QTRAP 5500 triple-quadrupole (Farmington, MA, USA) mass spectrometer was used to perform the metabolomic assays.

### Statistical Analysis

Statistical analyses were performed using Prism 9.0 (GraphPad Software). Normality of data was determined using the Shapiro-Wilk test. If data were normally distributed and there was equal variance as determined using the Brown-Forsythe test, 1-way analysis of variance with Tukey’s multiple-comparisons test was performed. If there was unequal variance, Brown-Forsythe and Welch analysis of variance with Dunnett multiple-comparisons test was completed. If the data were not normally distributed, the Kruskal-Wallis test and Dunn’s multiple-comparisons test were used when comparing two groups and the Mann Whitney U test for comparing two groups. Graphs display the mean or median value and all individual values. *p*-values of <0.05 were considered to indicate statistical significance. Hierarchical cluster analyses, sparse least discriminate analysis, random forest classification, and correlational heatmapping were performed using MetaboAnalyst software (https://www.metaboanalyst.ca/). The relative abundance of each protein was determined using Proteome Discover Software. Transcripts and proteins that were significantly correlated (*p*<0.05) with right ventricular systolic pressure and end-arterial elastance (Ea) were processed for Kyoto Encyclopedia of Genes and Genomes (KEGG) using ShinyGO 0.76.3 (http://bioinformatics.sdstate.edu/go/).

For detailed Materials and Methods, please refer to the Online Supplement.

## Results

### Ferroptosis Inhibition Mitigated Pulmonary Vascular Remodeling and Improved Right Ventricular Function in Monocrotaline Rats

We first evaluated the physiological effects of ferroptosis in monocrotaline (MCT) rats using a translational approach by starting treatment two weeks after MCT injection. Ferrostatin-1 (1 mg/kg) imparted important therapeutic effects as it increased pulmonary artery acceleration time, reduced right ventricular systolic pressure (RVSP) and effective arterial elastance (Ea), and suppressed small pulmonary arterial remodeling (**Figure 1A-D**). Accordingly, right ventricular (RV) function as defined by Ees/Ea, tricuspid annular plane systolic excursion (TAPSE), and RV free wall thickening were all improved with ferrostatin-1 (**Figure 1E-G**).

**Figure 1:**
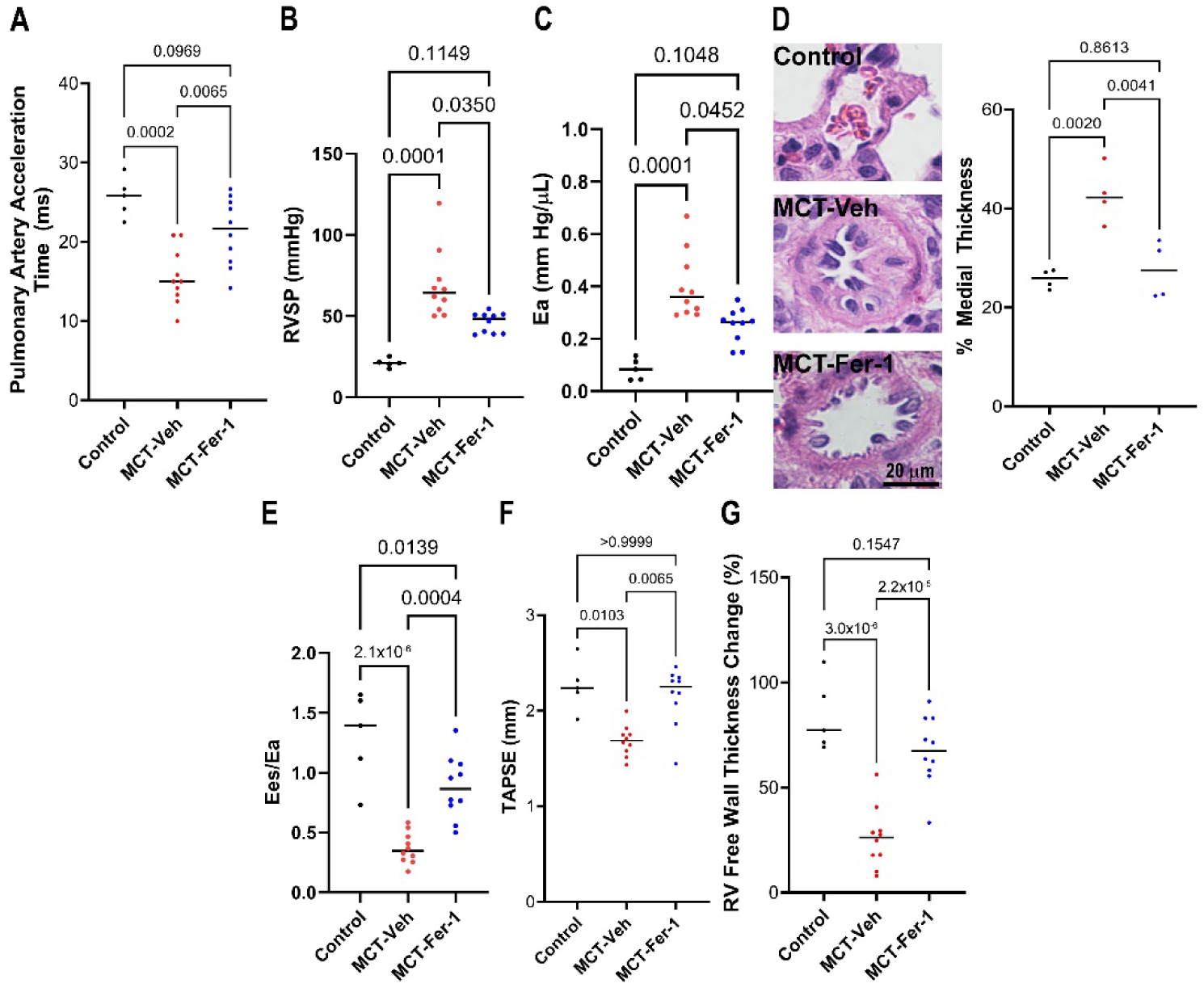
Ferrostatin-1 Treatment Combatted Pulmonary Hypertension Severity and Improved Right Ventricular Function in Monocrotaline Rats. (A) Ferrostatin-1 (1 mg/kg) treatment increased pulmonary artery acceleration time in MCT rats [Con: 26±1 (*n*=5), MCT-Veh: 16±1 (*n*=10), MCT-Fer-1: 21±1 ms (*n*=10)]. (B) Ferroptosis inhibition with ferrostatin-1 reduced right ventricular systolic pressure (RVSP) (Con: 21±1, MCT-Veh: 69±7, MCT-Fer-1: 46±2 mmHg), *p*-values determined by Kruskal-Wallis test. (C) Effective arterial elastance (E_a_) was lowered by ferroptosis inhibition (Con: 0.08±0.02, MCT-Veh: 0.40±0.04, MCT-Fer-1: 0.25±0.02 mmHg/μL), *p*-values determined by Kruskal-Wallis test. (D) Small vessel pulmonary arterial remodeling was blunted by ferrostatin-1 (Median values averaged from 4 animals per group: Con: 26±1, MCT-Veh: 43±3 MCT-Fer-1: 28±3, *p*-values determined by ANOVA with Tukey’s multiple comparisons test]. (E) Ferrostatin-1 treatment improved right ventricular-pulmonary artery coupling (E_es_/E_a_) (Con: 1.3±0.2, MCT-Veh: 0.37±0.04, MCT-Fer-1: 0.88±0.08), *p*-values determined by Brown-Forsythe and Welch ANOVA test), TAPSE (Con: 2.3±0.1, MCT-Veh: 1.7±0.05, MCT-Fer-1: 2.2±0.1, *p*-values determined by Kruskal-Wallis test) (F) and RV free wall thickening (Con: 84±8, MCT-Veh: 26±5, MCT-Fer-1: 67±5, *p*-values determined by ANOVA with Tukey’s multiple comparisons test) (G).

The summation of these data showed ferroptosis inhibition combatted pulmonary vascular remodeling and augmented RV function.

### RNA-Sequencing and Lung Mitochondrial Proteomics Analyses Identified Links Between Ferroptosis and Pulmonary Hypertension Severity

Next, we used RNA-sequencing (RNA-seq) and quantitative proteomics analyses of lung samples to perform an unbiased assessment of pathways that were dysregulated as PAH severity increased. RNA-seq data revealed ferroptosis was the second most enriched pathway associated with higher Ea (**Figure 2A**). Correlational heatmapping identified multiple positive and significant associations between ferroptotic transcripts and Ea (**Figure 2B and C**). When evaluating pathways induced by higher RVSP, ferroptosis was not identified in our unbiased assessment (**Supplemental Figure 1**). However, correlational heatmapping again identified multiple ferroptotic transcripts were positively associated with RVSP (**Supplemental Figure 1**).

**Figure 2:**
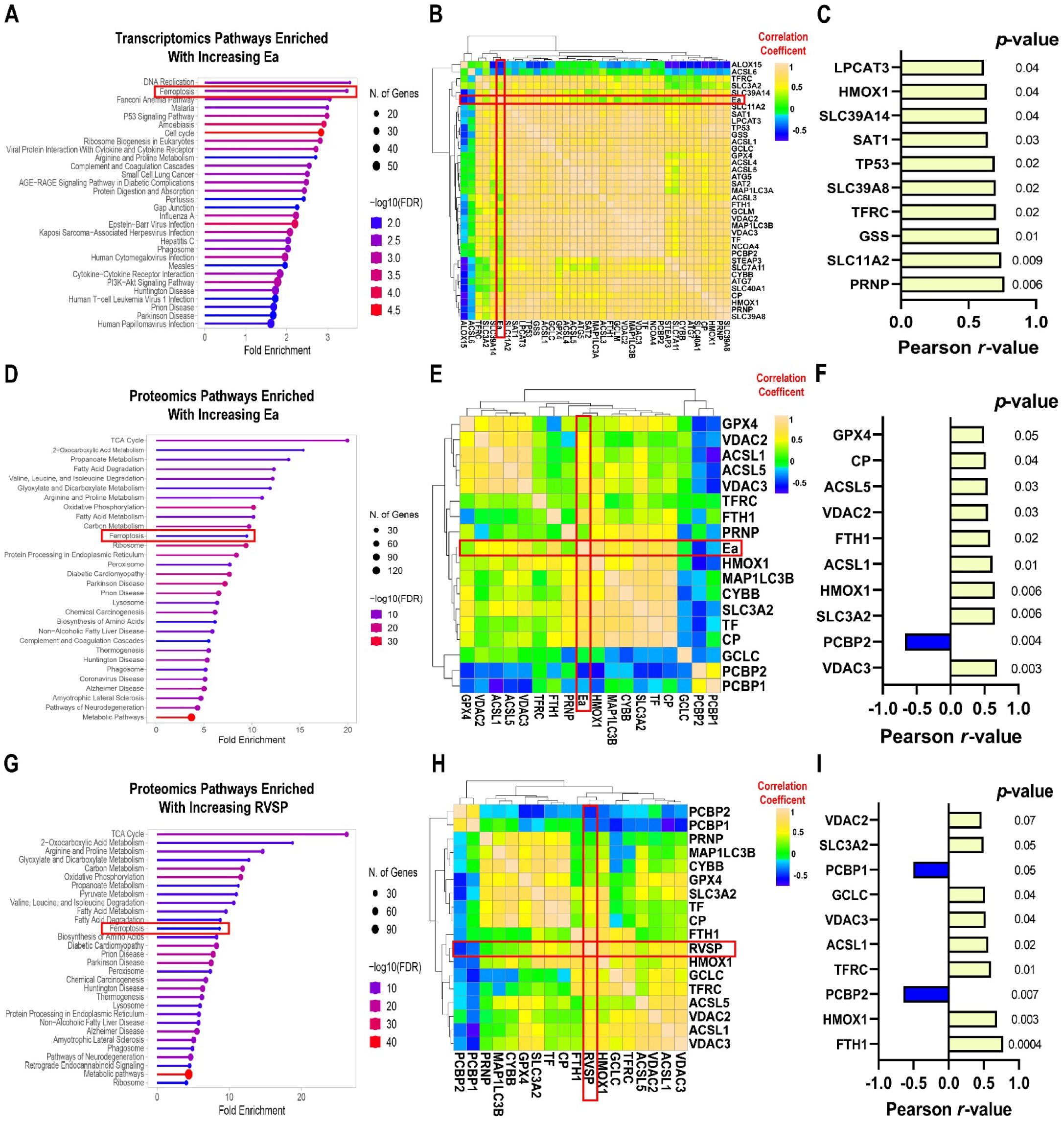
Lung RNA-sequencing and Proteomics Analyses Identified Associations Between Ferroptosis and Pulmonary Hypertension Severity. (A) Pathway analysis of transcripts isolated from whole lung specimens from (*n*=4 control, *n*=4 MCT-Veh, and *n*=3 MCT-Fer-1) significantly associated with Ea. (B) Correlational heat mapping of ferroptosis pathway transcripts and Ea. (C) Ten transcripts most strongly associated with Ea. (D) Pathway analysis of proteins (*n*=5 control, *n*=5 MCT-Veh, and *n*=6 MCT-Fer-1 animals) significantly associated with Ea. (E) Correlational heat map of ferroptosis proteins with Ea. (F) Ferroptosis proteins most strongly associated with Ea. (G) Pathway analysis of proteins significantly associated with right ventricular systolic pressure (RVSP). (H) Correlational heat map of ferroptosis proteins and RVSP. (I) Ten most strongly associated ferroptosis proteins with RVSP.

In our proteomics analysis, multiple metabolic pathways including the tricyclic acid cycle, fatty acid metabolism, oxidative phosphorylation, and amino acid metabolism were dysregulated as PAH severity increased. In agreement with our transcriptomic data, the ferroptosis pathway was enriched with increasing Ea and RVSP (**Figure 2 D and G**). Correlational heatmapping identified relationships between multiple ferroptotic proteins and Ea and RVSP (**Figure 2 E and H**). In particular, there were positive correlations between the pro-ferroptotic proteins, voltage-dependent anion-selective channel proteins 2 and 3 (VDAC 2/3), heme oxygenase 1 (HMOX1), and acyl-CoA synthetase long chain family members 1 and 5 (ACSL1/5) and Ea and RVSP (**Figure 2 F and I**). Conversely, the anti-ferroptotic protein poly(rC)-binding protein 2 (PCBP2) was negatively associated with Ea and RVSP (**Figure 2 F and I)**.

### Ferrostatin-1 Restructured the Lung Lipidomic Signature

Because our proteomics analysis identified several metabolic pathways were associated with pulmonary hypertension severity, we performed a metabolomics/lipidomics analysis of whole lung extracts. Visual examination of hierarchical cluster analysis and sparse least squares discriminate analysis demonstrated differences in the overall metabolomics/lipidomic signature when the three experimental groups were compared (**Supplemental Figure 2A and B**). Random forest classification identified the 15 metabolites that differentiated the three groups, which were predominantly lipid species and the amino acid, glycine (**Supplemental Figure 2C**). We subsequently used hierarchical cluster analysis to evaluate ferrostatin-1’s effects on three major lipid classes: phosphatidylcholines and lysophosphatidylcholines (PC, LPC), phosphatidylethanolamine and lysophosphatidylethanolamine (PE, LPE), and phosphatidylserine and lysophosphatidylserine (PS and LPS) as these lipids had the most significant differences when the three groups were compared. The most dramatic effects of ferrostatin-1 were on PC and LPC and PE and LPE. Ferrostatin-1 increased multiple PC and LPC species but reduced nearly all PE and LPE (**Supplemental Figure 2 D and E**). The effects of ferrostatin-1 on PS/LPS were less robust (**Supplemental Figure 2F**). When profiling amino acids, we found all amino acids other than glycine were elevated in MCT-Veh lungs when compared to controls, which was mostly mirrored in the ferrostatin-1 treatment animals with the exception of valine (**Supplemental Figure 2G**). Thus, ferroptosis inhibition appeared to have the greatest effect on lung lipid metabolism.

### Ferroptosis Inhibition Suppressed Classical Complement Activation and Cytokine Signaling

Next, we probed our ferroptosis-complement hypothesis by interrogating the coagulation/complement pathway using RNA-seq, proteomics, and immunofluorescence approaches. Hierarchical cluster analysis of RNA-seq data revealed ferrostatin-1 treatment blunted the upregulation of several coagulation/complement transcripts (**Figure 3A**). Correlational heatmapping displayed significant associations between multiple complement transcripts and RVSP and Ea (**Figure 3 B-E**). In particular, the transcript abundance of C1QA and C1QB were both correlated with Ea and RVSP (**Figure 3 C and E**). When this pathway was evaluated in our proteomic data, we also found ferroptosis inhibition prevented the upregulation of multiple complement proteins (**Figure 3F**). Correlational heatmapping again revealed several classical complement pathway proteins were associated with RVSP and Ea (**Figure 3 G-J**). To understand how ferrostatin-1 modulated complement activation in the pulmonary vasculature specifically, we used confocal microscopy to visualize perivascular complement deposition. In agreement with our proteomics and transcriptomics data, perivascular complement accumulation in MCT-Veh lungs was significantly elevated as compared to controls, but this response was mitigated by ferrostatin-1 (MCT-Veh: 1.8±0.2 fold increase compared to control, MCT-Fer-1: 1.1±0.2 fold increase compared to control *p*=0.02) (**Figure 3 K and L**).

**Figure 3:**
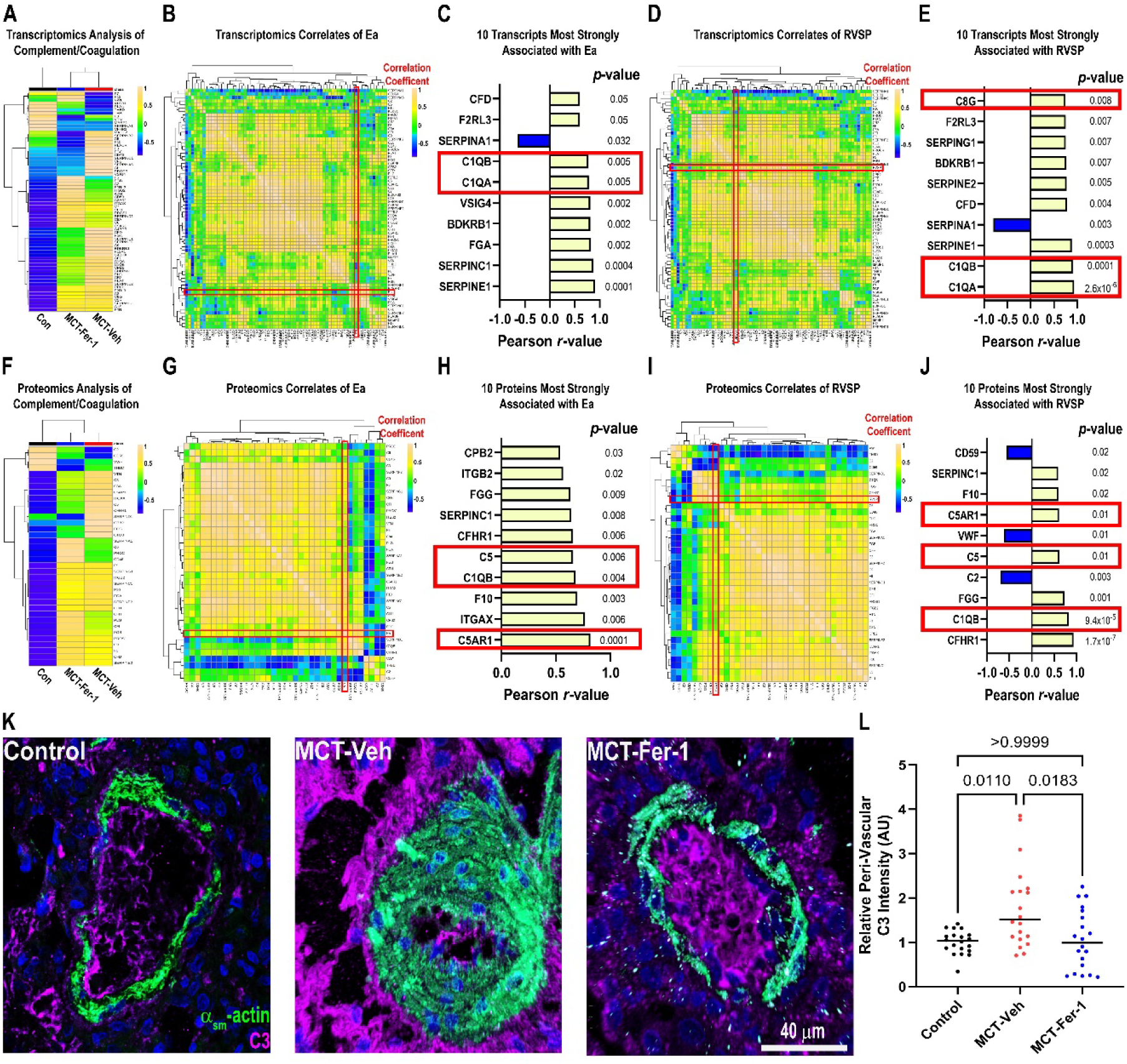
Ferroptosis Inhibition Mitigated Ectopic Complement Activation in the Lungs of MCT Rats. (A) Hierarchical cluster analysis of transcripts in the complement coagulation pathway. (A) Ferrostatin-1 reduced levels of many transcripts as compared to MCT-Veh. (B) Correlational heat map of complement/coagulation transcripts with Ea. (C) Identification of 10 transcripts most strongly associated with Ea. Classical complement components C1Qa and C1Qb were higher as Ea increased. (D) Correlational heat map of complement/coagulation transcripts with RVSP. (E) Ten transcripts most strongly associated with RVSP. (F) Hierarchical cluster analysis of proteins in the complement/coagulation cascade. (G) Correlational heat map of complement/coagulation proteins with Ea. (H) Identification of 10 proteins most strongly associated with Ea. (I) Correlational heat map of complement/coagulation proteins with RVSP. (J) Identification of 10 proteins most strongly associated with RVSP. (K) Representative confocal micrographs showed increased complement deposition around the pulmonary vasculature in MCT-Vehicle, which was mitigated by ferrostatin-1. (L) Quantification of perivascular C3 intensity. *p*-values determined by Kruskal-Wallis test and Dunn’s multiple comparisons test.

Because complement promotes cytokine and chemokine release, we evaluated how ferrostatin-1 impacted the lung inflammatory/cytokine signaling pathway in our RNA-seq data. Hierarchical cluster analysis showed ferrostatin-1 reduced the abundance of multiple transcripts in the inflammatory/cytokine pathway as compared to MCT-Veh (**Supplemental Figure 3A)** In particular, ferrostatin-1 reduced transcript abundance of nodal growth differentiation factor (NODAL), a secreted ligand of the transforming growth factor-beta superfamily of proteins, the transcript that exhibited the strongest relationship with Ea (**Supplemental Figure 3B**). In addition, ferrostatin-1 lowered transcript levels of tumor necrosis factor receptor superfamily member 11a (TNFRSF11A), a member of the tumor necrosis factor family whose transcript levels were associated with PAH severity (**Supplemental Figure 3B**). Finally, levels of pro-inflammatory chemokines including C-X-C motif chemokine ligands (CXCL) 6/10 and C-C motif chemokine ligands (CCL) 9/21 were reduced with ferrostatin-1 (**Supplemental Figure 3B**). Importantly, CXCL10 and CCL21 levels were positively associated with Ea (**Supplemental Figure 3C**).

### Deconvolution RNA-seq Analysis Revealed Ferrostatin-1 Modulated Endothelial Cell, Smooth Muscle Cell, Interstitial Macrophage Abundance and Metabolic and Inflammatory Phenotypes

To define how ferrostatin-1 modulated the cellular landscape of the pulmonary vasculature, we performed deconvolution RNA-seq analysis to define the changes in cell populations and pathways that were impacted by ferroptosis inhibition. All three experimental groups exhibited distinct cellular compositions in the lungs, but heterogeneity was observed (**Figure 4A, Supplemental Table 1**). Correlational heatmapping identified relationships between the proportion of cell types in the samples and Ea and RVSP (**Figure 4B and C**). Cell types known to underlie PAH pathobiology including smooth muscle cells, endothelial cells, and interstitial macrophages were most strongly associated with Ea and RVSP. Smooth muscle cells had the most robust correlational relationship with RVSP and Ea (**Figure 4D**). Importantly, ferrostatin-1 treatment partially mitigated the increase in lung smooth muscle cells when compared to MCT-Veh rats (**Figure 4D**). Additionally, ferroptosis inhibition partially suppressed the reduction of endothelial cells in the EA1 population and the increase in EA2 population. However, the reduction in EA1 as opposed to the increase in EA2 cells (data not shown) was more significantly associated with PAH severity (**Figure 4E and F**). Additionally, greater abundance of interstitial macrophages was associated with more severe PAH as MCT-Veh rats had the highest numbers of interstitial macrophages (**Figure 4 E and F**). Ferroptosis inhibition lowered interstitial macrophage numbers as compared to MCT-Veh (**Figure 4 E and F**). Pathway analysis delineated how ferrostatin-1 modulated the transcriptional environment in highly abundant cells. Ferroptosis inhibition suppressed the activation of multiple pathways in both endothelial cells and macrophages. In both cell types, changes in the DNA damage, cellular replication (PI3K/AKT/mTOR signaling and mitotic spindle), heme metabolism, and lipid handling (adipogenesis) pathways were observed (**Figure 4G**). Finally, we used confocal microscopy to evaluate how ferroptosis modulated macrophage infiltration into the pulmonary vasculature. There was a significant increase in perivascular CD11b^+^ cells in the MCT-Veh animals as compared to control animals, which ferrostatin-1 treatment reduced (MCT-Veh: 10.4±0.9 CD11b^+^ cells/arteriole, MCT-Fer-1: 4.3±0.5 CD11b^+^ cells/arteriole *p*=2.5×10^-5^) (**Figure 4H**).

**Figure 4:**
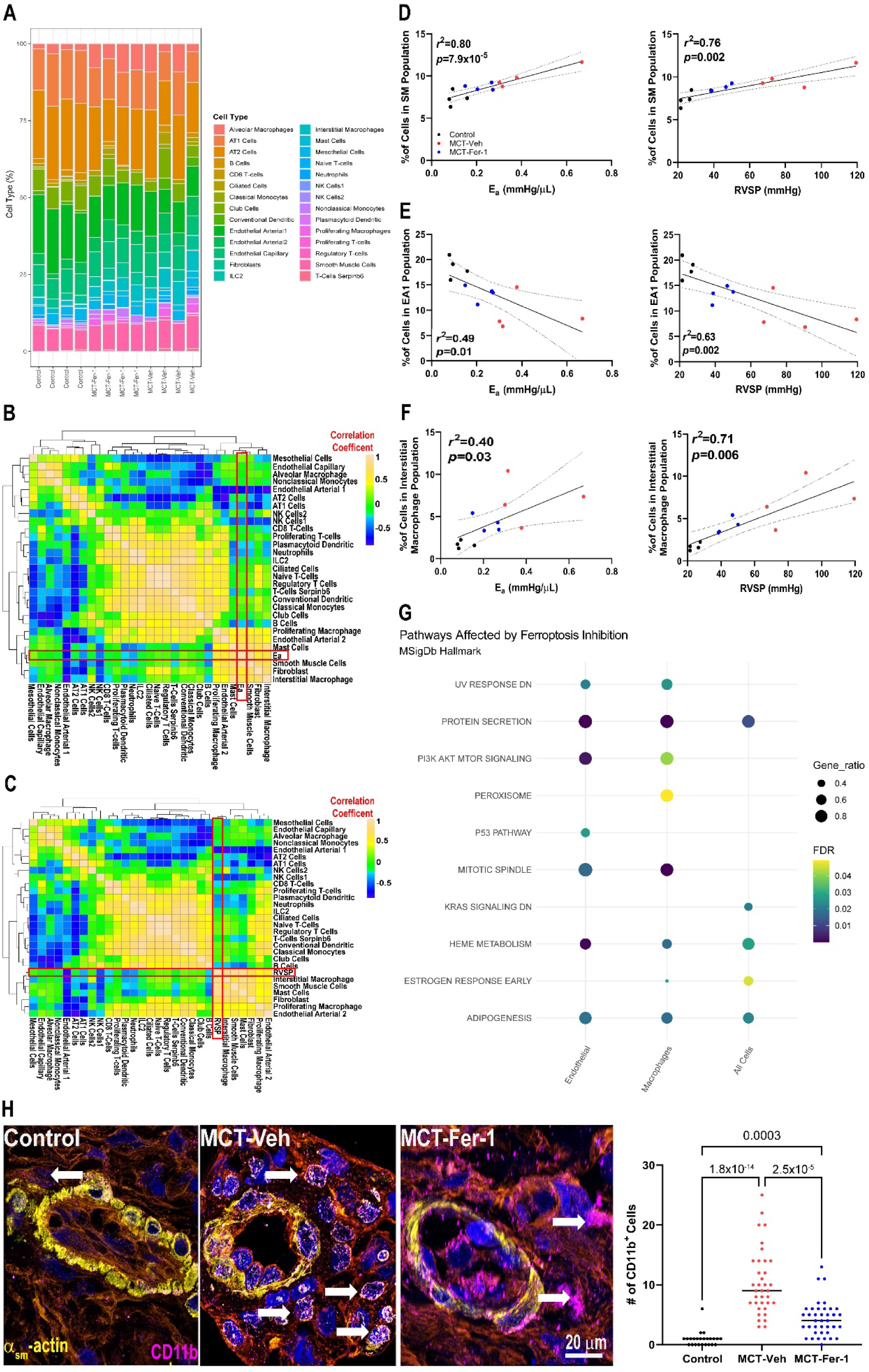
Deconvolution RNA-sequencing Demonstrated Ferroptosis Inhibition Modulated Endothelial, Smooth Muscle, and Interstitial Macrophage Abundances and Gene Activation Patterns. (A) Map of cell type abundances in control, MCT-Veh, and MCT-Fer-1 animals as determined by deconvolution RNA-seq (*n*=4 control, *n*=4 MCT-Veh, and *n*=4 MCT-Fer-1). Correlational heat mapping identified relationships between cell type abundances and Ea (B) and RVSP (C). (D) Relationships between smooth muscle cell abundance and Ea and RVSP. (E) Relationships between endothelial cell population 1 (EA1) and Ea and RVSP. (F) Relationships between interstitial macrophage population and Ea and RVSP. (G) Pathway analysis of transcriptomic changes in endothelial cells, macrophages, and total cells with ferrostatin-1 treatment. (H) Representative images of pulmonary arteriole demonstrating perivascular CD11b^+^ positive cells (arrow) and quantification of peri-vascular CD11b^+^ cells. *p*-values determined by Kruskal-Wallis test and Dunn’s multiple comparisons test.

### Ferroptotic Media from Human Pulmonary Artery Endothelial Cells Promoted Pathogenic Changes in Human Pulmonary Artery Smooth Muscle Cells and Human Monocytes

To supplement our multi-omics analysis and provide more mechanistic data using human cells, we next determined how media from ferroptotic human PAEC, which contained ferroptotic DAMPs, impacted *in vitro* phenotypes of human PASMC and human monocytes. Incubating PASMC with ferroptotic PAEC DAMPs (**Figure 5A**) altered the transcriptomic landscape as there were 699 differentially expressed genes when PASMC treated with ferroptotic media were compared to PASMC treated with control media. Hierarchical cluster analysis demonstrated the two conditions resulted in distinct RNA signatures (**Figure 5B**). Pathway analysis of transcripts upregulated by ferroptotic media suggested there were alterations in metabolic pathways including amino acid metabolism, hypoxia inducible factor-1 (HIF1), and mammalian target of rapamycin (mTOR) signaling (**Figure 5C**). Pathway analysis of the downregulated transcripts identified an enrichment of pathways important for regulation DNA integrity/repair and cell cycle regulation (**Figure 5C**). Next, we used confocal microscopy as a cellular readout of our RNA-seq results. Ferroptotic media disrupted the mitochondrial network as characterized by increased fragmentation and a reduced mitochondrial footprint (**Figure 5D**). Because PASMC DNA repair and cell cycle regulation gene programs were altered by ferroptotic PAEC DAMPs, we evaluated PASMC proliferation status because DNA damage is associated with PASMC replication in PAH^18^. Ferroptotic DAMPs heightened PASMC proliferation as the proportion of Ki67^+^ cells was significantly higher than control media (Control Media:40±5%, Ferroptotic Media: 67±6%, *p*=0.03) (**Figure 5E**)

**Figure 5:**
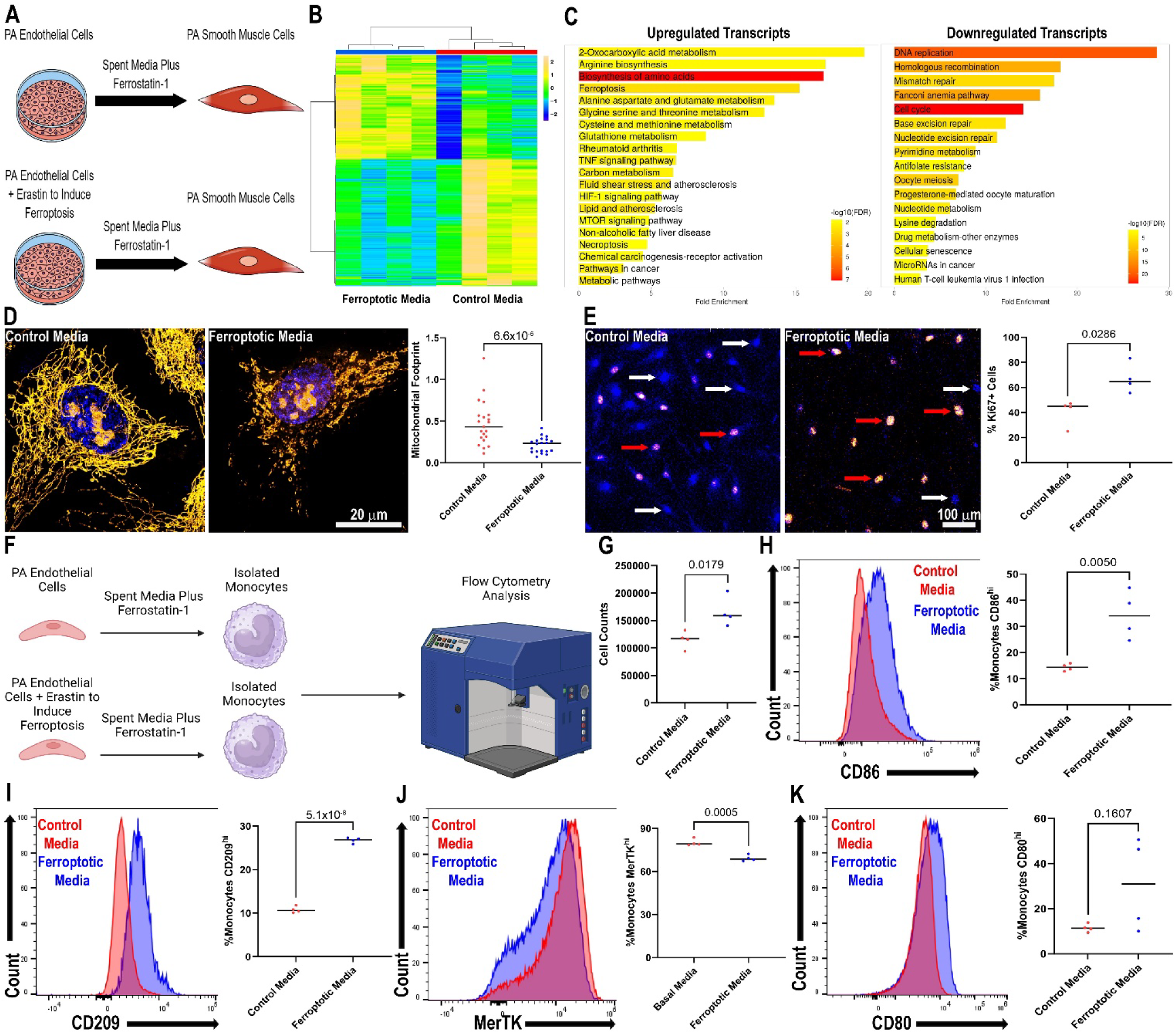
DAMPs from Ferroptotic Human Pulmonary Arterial Endothelial Cells Recapitulated Pathogenic Changes in Human Pulmonary Artery Smooth Muscle Cells and Promoted a Pro-Inflammatory Phenotype in Human Monocytes *In Vitro*. (A) Diagram of experimental approach to evaluate the effects of ferroptotic pulmonary artery endothelial cells on pulmonary artery smooth muscle cell phenotypes *in vitro*. (B) Hierarchical cluster analysis of top 699 transcripts from RNA isolated from *n*=4 replicates of PASMC treated with control media and *n*=4 replicates of PASMC treated with ferroptotic media. (C) KEGG pathway analysis of upregulated and downregulated transcripts comparing control and ferroptotic media treated PASMC. (D) Super resolution confocal micrographs of PASMC incubated with control and ferroptotic PAEC spent media stained with MitoTracker Orange and DAPI and quantification of the mitochondrial footprint. *p*-value determined by Mann Whitney test. (E) Confocal micrographs of PASMC stained with DAPI and Ki67 to identify replicating cells. Red arrows indicate Ki67^+^ cells, white arrows indicate Ki67^-^ cells. Quantification of the proportion of Ki67^+^ cells, *p*-value determined by Mann Whitney test. (F) Diagram of experimental conditions employed to evaluate *in vitro* effects of ferroptotic PAECs on isolated human monocytes. (G) Quantification of the number of viable monocytes following media exposure. *p*-value determined by unpaired *t*-test. Flow cytometry analysis of monocyte expression of CD86 (H), CD209 (H), MerTK (J), and CD80 (K). *p*-values determined by unpaired *t*-test in H, I, and J and unpaired *t*-test with Welch correction in K.

Then, we used flow cytometry to probe how PAEC ferroptotic DAMPs altered monocyte inflammatory status *in vitro* (**Figure 5F and G**). We first found ferroptotic media promoted monocyte survival (**Figure 5H**). Moreover, ferroptotic media induced a pro-inflammatory signature, as there were significantly increased proportions of cells that expressed high levels of the pro-inflammatory markers CD86 (Control Media: 14±1%, Ferroptotic Media: 34±5%, *p*=0.005), CD209 (Control Media: 11±0.4%, Ferroptotic Media: 37±0.3%, *p*=5.1×10^-8^), a significantly lower proportion of cells that expressed the anti-inflammatory and pro-efferocytic marker MerTK (Control Media: 80±1%, Ferroptotic Media: 69±1%, *p*=0.0005), and a nonsignificant increase in the proportion of cells expressing high levels of CD80 (**Figure 5G-K**). These data suggested ferroptotic PAEC DAMPs stimulated survival and a pro-inflammatory phenotype in human monocytes.

### Viral-Mediated Endothelial Cell Overexpression of the Pro-Ferroptotic Protein ACSL4 Induced an Inflammatory Pulmonary Hypertension Phenotype

Next, we used a genetic approach to complement our small-molecule data to evaluate the role of ferroptosis in rodent pulmonary hypertension by comparing the effects of intratracheal delivery of AAV1-*Acsl4* (*n*=5) or AAV1-GFP (*n*=5), driven by the endothelial cell specific cadherin-5^19^ promoter, in rats exposed to low dose (30 mg/kg) monocrotaline (**Figure 6A**). Immunofluorescence analysis demonstrated heightened ACSL4 immunoreactivity in endothelial cell layer adjacent to PASMC in the AAV1-*Acls4* treated rats as compared to AAV1-GFP rats (**Figure 6A**). Physiological characterization revealed AAV1-*Acsl4* induced moderate pulmonary hypertension with significantly reduced pulmonary artery acceleration time (AAV1-GFP: 19±1 ms, AAV1-*Acsl4*: 15±1 ms, *p*=0.02) (**Figure 6B**) and elevated RVSP (AAV1-GFP: 33±2 mmHg, AAV1-*Acsl4*: 55±3 mm Hg, *p*=0.0004) (**Figure 6C**) and Ea (AAV1-GFP: 0.1±0.02 mmHg/μL, AAV1-*Acsl4*: 0.3±0.05 mmHg/μL, *p*=0.0004) (**Figure 6D**). Unfortunately, one of the AAV1-*Acsl4* rats died during the hemodynamic study so only 4 animals had complete hemodynamic characterization. AAV1-*Acsl4* heightened pathological small vessel pulmonary vascular remodeling (**Figure 6E**) and peri-vascular complement deposition (**Figure 6F**). The alterations in the pulmonary vasculature induced right ventricular cardiomyocyte hypertrophy (**Figure 6G**) and compromised right ventricular function (**Figure 6 H-J**). In conclusion, these data demonstrated genetically-initiated PAEC ferroptosis via ACSL4 overexpression induced an inflammatory pulmonary hypertension.

**Figure 6:**
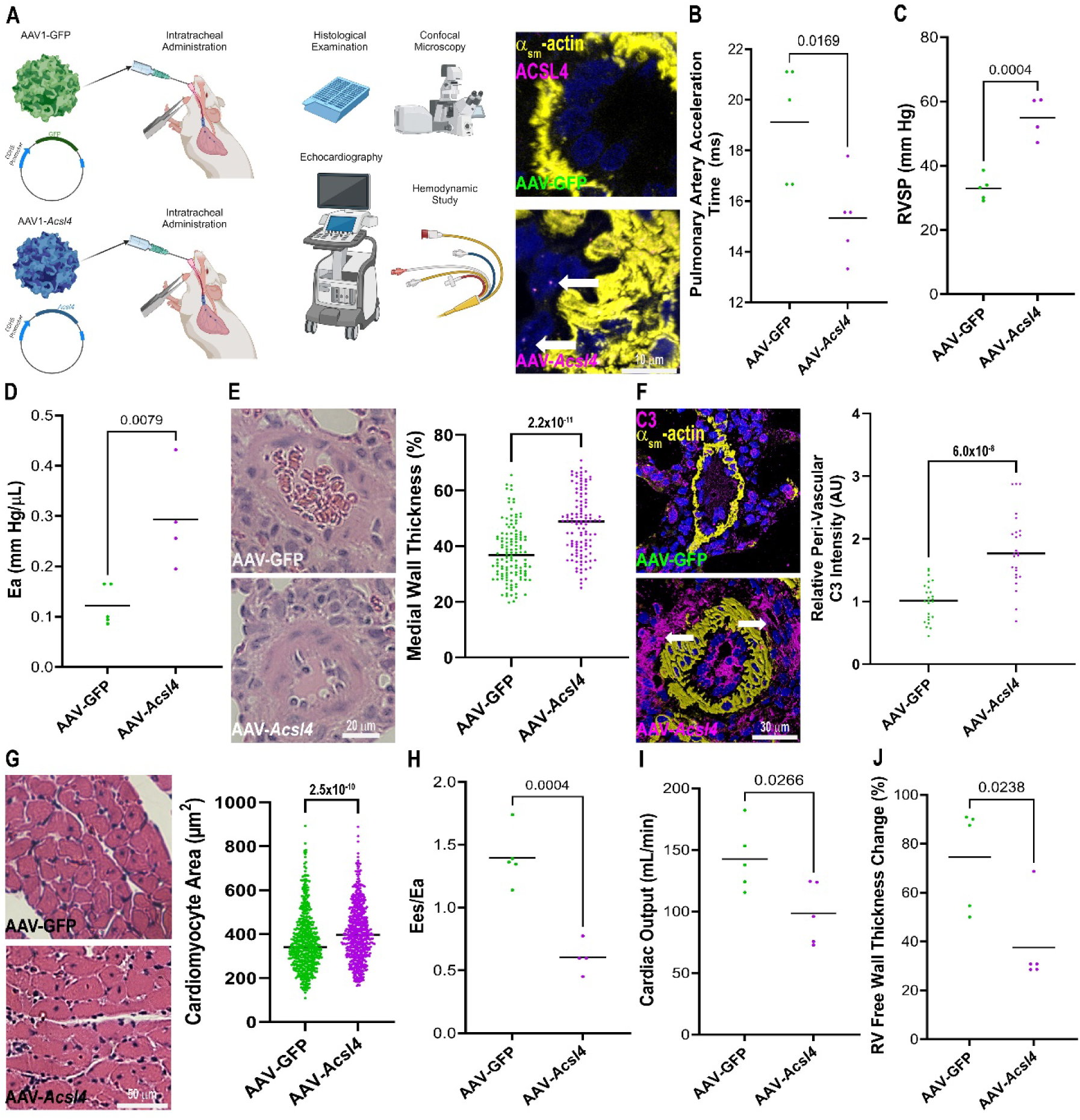
AAV1-Mediated Endothelial *ACSL4* Overexpression Induced Perivascular Complement Deposition, Pulmonary Hypertension, and Right Ventricular Dysfunction in Rats. (A) Schematic presentation of experimental approach of rats treated with 0.3×10^11^ vector genomes of AAV1-*Acsl4* or 0.3×10^11^ vector genomes of AAV1-GFP with concurrent exposure to low dose (30 mg/kg) monocrotaline with confocal micrographs demonstrating heightened ACSL4 immunoreactivity in the endothelial cell layer (arrows). AAV1-*Acsl4* treatment induced pulmonary hypertension as pulmonary artery acceleration time was reduced [AAV1-GFP: 19±1 ms (*n*=5) AAV1-*Acsl4*: 15±0.7 ms (*n*=5)] (B) and RVSP [AAV1-GFP: 33±2 mm Hg (*n*=5), AAV1-*Acsl4*: 55±3 mm Hg (*n*=4)] (C), Ea [AAV1-GFP: 0.12±0.2 (*n*=5), AAV1-*Acsl4*: 0.29±0.05 (*n*=4)] (D) were increased and small vessel remodeling was heightened (E). (F) Representative confocal micrographs and quantification of C3 immunofluorescence intensity demonstrated AAV1-*Acsl4* promoted perivascular complement deposition. AAV1-*Acsl4* induced RV cardiomyocyte hypertrophy (AAV1-GFP: 364±5 μm^2^, AAV1-*Acsl4*: 407±5 μm^2^) (G) and RV dysfunction as RV-PA coupling was compromised [AAV1-GFP: 1.4±0.1 (*n*=5), AAV1-*Acsl4*: 0.6±0.1 (*n*=4)] (H) and cardiac output [AAV1-GFP: 140±12 mL/min (*n*=5) (I), AAV1-*Acsl4*: 99±11 mL/min (*n*=5)] and RV free wall thickening [AAV1-GFP: 75±9% (*n*=5), AAV1-*Acsl4*: 38±8% (*n*=5)] were impaired (J). *p*-values as determined by unpaired t-test (B, C, H, and I) or Mann-Whitney test (D-G, and J)

### Presence of Single-Nucleotide Polymorphisms in Ferroptosis Genes Potentially Associated with More Severe Pulmonary Hypertension

Finally, we evaluated how the presence of single-nucleotide polymorphisms (SNPs) in ferroptosis genes impacted noninvasive and invasive measures of pulmonary hypertension in a mixed cohort of patients evaluated at the Vanderbilt BioVU biorepository. The presence of six SNPs in ferroptosis genes, which were characterized as variants of unknown significance with four SNPs resulting in amino acid substitutions and two SNPs were located in intronic sequences, were associated with more severe pulmonary hypertension using both echocardiographic and hemodynamic measures, however the overall frequency was low (**Supplemental Table 2**). Therefore, we pooled all six SNPs and compared how carriers of ferroptosis SNPs compared to those lacking potentially pathogenic SNPs. In an unadjusted analysis, carriers had significantly higher RVSP [Carriers: 63.9 (45.0-80.6), Noncarriers: 46.8 mmHg (35.0-62.0), *p*=0.0001] as determined by echocardiography (**Figure 7A**). Furthermore, invasively measured mean pulmonary arterial pressure (mPAP) [Carriers: 40.0 (25.0-51.0), Noncarriers: 31.0 mmHg (22.0-41.0), *p*=0.02], and pulmonary vascular resistance (PVR) [Carriers: 4.3 (2.4-6.9), Noncarriers: 2.6 Wood units (1.7-4.6), *p*=0.001] were higher in those harboring ferroptosis SNPs (**Figure 7 B-C**). Because this was a mixed cohort, we performed a multivariate regression analysis and even after correcting for age, race, sex, presence of heart failure, chronic obstructive pulmonary disease, and use of pulmonary vasodilators, ferroptotic SNPs were independently associated with higher RVSP, mPAP, and PVR (**Figure 7 D-E**). These data highlight a potential link between ferroptosis and pulmonary hypertension severity in humans. However, future studies are needed to determine how these SNPs regulate proteins function or alter splicing to provide further understanding of how these SNPs may impact ferroptotic regulation.

**Figure 7:**
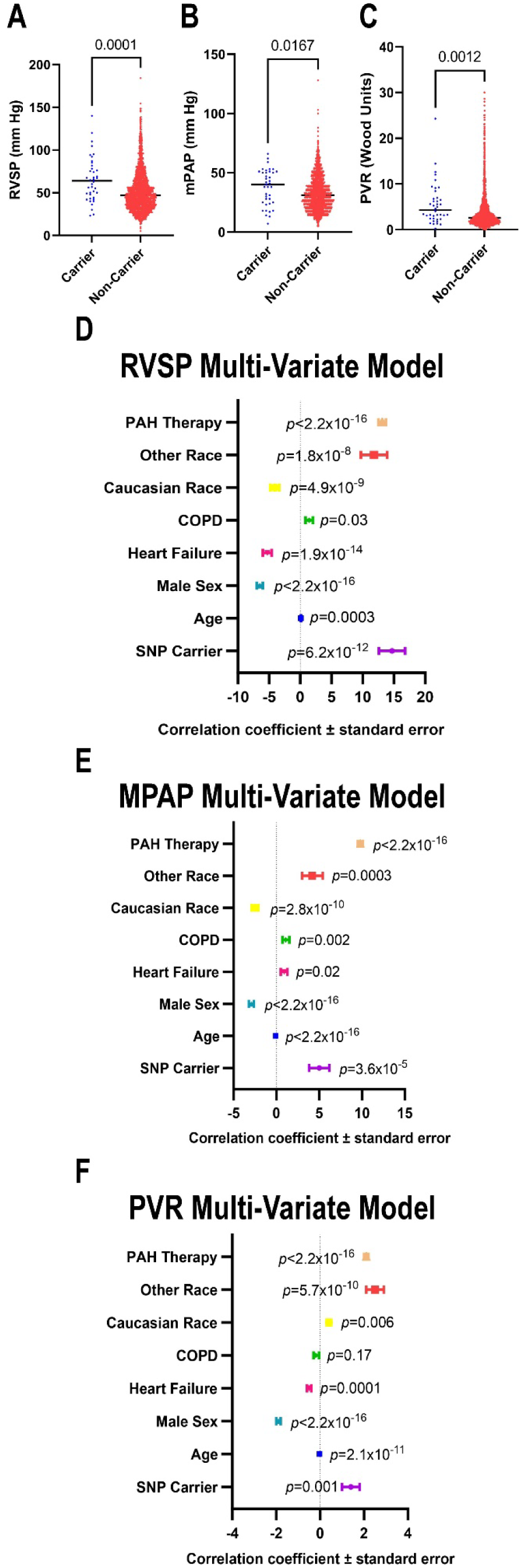
Presence of Single Nucleotide Polymorphisms in Ferroptosis Regulating Genes Was Associated With More Severe Pulmonary Hypertension. (A) Patients harboring SNPs in ferroptosis genes had higher right ventricular systolic pressures [Carriers: 63.9 (45.0-80.6), Noncarriers 46.8 (35.0-62.0), median (25^th^-75^th^ percentiles)] as determined by echocardiography, (B) invasively measured mean pulmonary arterial pressure [Carriers: 40.0 (25.0-51.0), Noncarriers 31.0 (22.0-41.0), median (25^th^-75^th^ percentiles)], and pulmonary vascular resistance [Carriers: 4.3 (2.4-6.9), Noncarriers 2.6 (1.7-4.6), median (25^th^-75^th^ percentiles)] as determined by invasive hemodynamic studies (C). *p*-values as determined by Mann-Whitney U-test. Forest plots depicted correlational co-efficient with standard errors and calculated *p*-values when RVSP (D), mPAP (E), and PVR (F) were modeled in multivariate regression analysis.

## Discussion

In summary, our translational rodent studies show small molecule inhibition of ferroptosis mitigates PAH and improves right ventricular function in MCT rats even when implemented after PAH has developed. Transcriptomics and proteomics analyses demonstrate a relationship between ferroptosis induction and PAH severity, which provides unbiased data to support our physiological findings. Our metabolomics/lipidomics experiments suggest ferrostatin-1 most robustly modulates lung lipid metabolism. In the pulmonary vasculature, suppression of ferroptosis alters endothelial and pulmonary artery smooth muscle cell numbers and gene activation patterns. Additionally, ferrostatin-1 mitigates pathological immune activation in the lung by suppressing perivascular complement deposition, lowering cytokines and chemokines, and reducing interstitial macrophage abundance. *In vitro* studies support our deconvolution RNA-seq experimental results as media containing DAMPs from ferroptotic PAEC promotes pathogenic changes in human PASMC marked by alterations in metabolic pathways including HIF/mTOR signaling and pathways responsible for DNA homeostasis. These transcriptomic changes are paired with mitochondrial morphological alterations and heightened proliferation rates. In addition, ferroptotic PAEC DAMPs induce a pro-inflammatory phenotype in human monocytes. Furthermore, we provide genetic evidence that PAEC ferroptosis is sufficient to induce pulmonary hypertension as AAV1-*Acsl4* results in an inflammatory pulmonary hypertension with associated right ventricular dysfunction in rats. Finally, a human genetic-association study reveals SNPs in ferroptosis genes are potentially associated with more severe pulmonary vascular disease burden. In conclusion, our data support a model in which ferroptotic PAECs release DAMPs that promote ectopic complement activation, serve as a signal to recruit and activate pro-inflammatory macrophages to the pulmonary vasculature, and alter the transcriptomic landscape of PASMC in a manner that recapitulates PAH pathobiology. The summation of these cellular and molecular alterations ultimately promote pulmonary hypertension (**Central Figure**). However, future work targeting each of these steps *in vivo* is required to further refine our working model.

Our data suggest PAEC ferroptosis drives pulmonary hypertension by both altering the PASMC genetic program and activating inflammatory monocytes/macrophages. Incubation of PASMC with ferroptotic endothelial cell media modulates pathways already implicated in PAH pathobiology. First, we show ferroptotic PAECs reduces PASMC DNA repair pathway transcript abundances. DNA damage triggers PASMC proliferation and dysfunction as there is increased DNA damage in PASMC in PAH^18^. Importantly, modulation of DNA damage repair pathways counteracts pulmonary vascular remodeling and reduces pulmonary hypertension severity in rodents^18,20^. Moreover, transcriptomic data suggest ferroptotic PAEC media activates HIF1^21^ and mTOR^22^, two pathways with a wealth of data demonstrating their pathogenic roles in PAH. In addition, ferroptotic endothelial cell DAMPs increase monocyte survival and promote a pro-inflammatory phenotype *in vitro*, demonstrating another established link to PAH pathobiology. These data are in line with other manuscripts showing inhibition of ferroptosis suppresses inflammatory signaling via the NOD-, LRR-, and pyrin domain-containing protein 3 (NLRP3) inflammasome activation in the lung^6^ and heightened levels of interstitial macrophage in human PAH lungs^13^. Certainly, future work is required to understand what specific components of the DAMPs released from ferroptotic PAECs drive the changes in PASMC and macrophages so more discrete and targetable approaches can be employed.

Our findings that ferroptosis induces both metabolic derangements and pathological inflammation in the lungs has potential to link two heavily implicated molecular phenotypes in PAH pathology to one targetable pathway. In PAH, derangements in lung metabolism including altered fatty acid oxidation^23,24^, heightened anaerobic glucose metabolism^23^, and disrupted amino acid utilization^25–27^ are well documented. Ferroptosis has the potential to impact several catabolic pathways via its ability to suppress electron transport chain activity^28^, a final common pathway for nearly all metabolic processes. While targeting specific metabolic arms including suppression of fatty acid oxidation^29^ and activation of glucose oxidative metabolism^30^ has robust therapeutic effects in preclinical PAH, the translation of these studies to humans has not been met with overwhelming success^31^. Perhaps, targeting one isolated branch of metabolism is insufficient to counteract human PAH pathogenesis, and more broad-based metabolic interventions would garner greater success. In addition, ferroptosis antagonism also combats ectopic complement activation, lowers levels of several cytokines/chemokines, and mitigates macrophage infiltration demonstrates the anti-inflammatory effects of ferrostatin-1 in the pulmonary vasculature. While there is strong data showing many individual cell types or cytokines/chemokines promote PAH in rodents^32^; unfortunately the translation of these findings to humans has not been as fruitful as desired. In particular, antagonism of interleukin-6 signaling with tocilizumab^33^ or depletion of B-cells with rituximab^34^ did not significantly change PAH severity in two small clinical trials. Perhaps suppressing an individual cytokine or immune cell type may not be the most effective therapy in humans as available data suggests there is widespread immune activation in PAH^32^. In fact, the cross-talk between immune and pulmonary vascular cells is crucial for PAH progression/severity as there is greater immune-pulmonary vascular interactions in hereditary PAH as compared to idiopathic PAH, and this finding correlates with hemodynamic severity^35^. Because our data suggests ferroptosis both dampens mitochondrial metabolism and initiates innate immune system activation in the pulmonary vasculature, perhaps approaches to counteract ferroptosis may lead to greater translational success by combatting both metabolic and inflammatory mechanisms.

We show ferroptosis inhibition mitigates pulmonary hypertension severity in MCT rats and ferroptosis induction via AAV-*Acsl4* causes pulmonary hypertension. These results are congruent with additional preclinical and human data that support our hypothesis that ferroptosis promotes pulmonary hypertension. First, several ferroptosis-suppressing molecules exhibit therapeutic efficacy in relevant preclinical models of pulmonary vascular disease including MCT, Sugen-hypoxia, and chronic hypoxia rats and lambs with persistent pulmonary hypertension of the newborn (**Table 1**). In addition, KEGG and WIKI pathway analysis of upregulated transcripts in PAH endothelial cells analyzed by single cell RNA-seq^36^ identifies ferroptosis as an enriched pathway in PAH (**Supplemental Figure 4**). Furthermore, a proteomic study demonstrates a 35% reduction in glutathione peroxidase-4, an anti-ferroptotic protein, in PAECs from PAH patients^37^. Moreover, there are higher systemic levels of lipid peroxidation in PAH patients as urinary abundance of isoprostaglandin F2a are significantly higher than controls^38^. Finally, circulating levels of transferrin receptor, a proposed biomarker of ferroptosis^39^, are elevated in PAH patients and can independently predict mortality^40^. In addition, higher circulating levels of transferrin receptor levels are associated with more severe pulmonary hypertension in patients with chronic lung disease^41^. Thus the summation of our results, other published pre-clinical therapeutic evaluations, and human transcriptomics, proteomics, and biomarker data suggest ferroptosis may be a targetable pathway to suppress adverse pulmonary vascular remodeling.

**Table 1:** Numerous Ferroptosis Suppressing Drugs Combat Pulmonary Hypertension in Multiple Preclinical Models.

| Small Molecules | Proposed Mechanisms of Ferroptosis Suppression | Animal Models With Therapeutic Effects |
| --- | --- | --- |
| AICAR and metformin | AMPK Activation | Chronic hypoxia-induced pulmonary hypertension in rats, Sugen-hypoxia induced pulmonary hypertension, monocrotaline rats<sup>48</sup> |
| Apocynin | Reduces reactive oxygen species required for lipid peroxidation in the NOX family of proteins | Lambs with persistent pulmonary hypertension of the newborn<sup>49</sup> |
| Deferoxamine | Iron chelation to prevent iron-dependent lipid peroxidation | Chronic hypoxia-induced pulmonary hypertension in rats<sup>50</sup> |
| Ferrostatin-1 | Suppresses ferroptosis directly and mitigates NLRP3 Activation in the lungs | Monocrotaline-induced pulmonary hypertension in rats<sup>6,8</sup> |
| MK866 | Inhibits lipoxygenase-mediated lipid peroxidation | Chronic hypoxia-induced pulmonary hypertension in rats<sup>51</sup> |
| Rosiglitazone/pioglitazone | Antagonizes ACSL4-induced conversion of free fatty acids into fatty CoA esters that then become peroxidated | Monocrotaline rats, sugen-hypoxia rats, chronic hypoxia rats, ApoE<sup>-/-</sup> mice, transgenic TGFβ1 mice<sup>52</sup> |
| Rapamycin | mTOR Suppression | Monocrotaline-induced pulmonary hypertension in rats, chronic hypoxia-induced pulmonary hypertension in mice and rats, Sugen-hypoxia rats<sup>53</sup> |
| Vildagliptin | Blocks DPP4-mediated lipid peroxidation | Monocrotaline-induced pulmonary hypertension in rats, chronic hypoxia-induced pulmonary hypertension in rats<sup>54</sup> |
| Zileuton | Inhibits lipoxygenase-mediated lipid peroxidation | Monocrotaline pulmonary hypertension<sup>55</sup> |

**Abbreviations:** AMPK: Adenosine monophosphate activate protein kinase, NOX: NADPH oxidases, ACSL4: acyl-CoA synthetase long-chain family member 4, mTOR: mammalian target of rapamycin, DPP4: dipeptidyl-dippeditase-4.

In our proteomics analysis of the lungs, we highlighted multiple mitochondrial metabolic pathways which are altered as pulmonary hypertension severity increases. However, the metabolomics/lipidomics analysis suggested the greatest effect of ferrostatin-1 was on lipid metabolism. In particular, ferrostatin-1 increases levels of multiple phosphatidylcholines (**Supplemental Figure 2**), which may have direct relevance to mitochondrial function because phosphatidylcholines are a major component of the mitochondrial membrane system^42^. Perhaps the greater concentrations of phosphatidylcholines contributes to the proteomic changes in other metabolic pathways via the stabilization and preservation of mitochondria. In addition, phosphatidylcholines are required for inner membrane protein translocases in the mitochondria^42^. A change in phosphatidylcholines could also alter mitochondrial protein stability as matrix and inner membrane proteins may not be properly trafficked into the mitochondria in the PAH lung due to the dysregulation of phosphatidylcholines. Additionally, ferrostatin-1 lowered the levels of nearly all phosphatidylethanolamines profiled (**Supplemental Figure 2**). This may be an important feedback mechanism to mitigate ferroptosis because peroxidation of phosphatidylethanolamines is crucial for ferroptosis initiation^43^. Surprisingly, ferrostatin-1 had almost no effect on lung amino acid abundances with the exception of reducing levels of valine, a branched chain amino acid. While amino acids regulate ferroptosis homeostasis in other conditions^44^, our data suggest amino acids may not play a dominant role in endothelial cell ferroptosis induction in PAH. In conclusion, the strongest effects of ferrostatin-1 on lung metabolism appears to be on lipid homeostasis, and these changes may underlie some of the beneficial physiological effects of ferroptosis antagonism.

Our deconvolution RNA-seq analysis identifies changes in pulmonary endothelial cell, smooth muscle cell, and interstitial macrophage number and gene activation patterns. However, other immune cell types undoubtedly contribute to pulmonary vascular remodeling, but our approach may have not been sensitive enough to detect changes in these lower abundant immune cell populations. Certainly, there is literature demonstrating T-cells^45,46^ and B-cells^47^ play important roles in pulmonary vascular remodeling, however we did not observe changes in these cell populations with ferroptosis inhibition (**Supplemental Table 1**). In addition, we did not see a profound alteration in mast cell numbers with ferrostatin-1 treatment (**Supplemental Table 1**). With the limitations of our approach, it appears ferroptosis inhibition exerted the greatest effects on endothelial cells, smooth muscle cells, and recruited macrophages. The complex interplay between these distinct cell types in the pulmonary vasculature appears to be responsible for adverse effects of ferroptosis in pathological pulmonary vascular remodeling (**Central Figure**).

## Limitations

Our work has important limitations that we must acknowledge. First, we used whole lung tissue in our proteomics and metabolomics analyses, and unfortunately defining what cell types are contributing to the observed changes cannot be determined. We tried to overcome some of these limitations by using deconvolution RNA-seq to nominate changes in cell numbers and gene activation patterns and by performing *in vitro* analyses of the most altered cell types: monocytes and PASMC. Finally, our SNP analysis was conducted in a mixed population of pulmonary hypertension patients in a small cohort. When we evaluated these SNPs in a larger secondary cohort: the UK PAH Cohort study^43^, we found they were in very low abundance and not associated with differences in PAH severity in an unadjusted analysis (**Supplemental Figure 5**). Finally, we evaluated phenotypes only in males because of the more robust phenotype in male rodents.

## Non-standard Abbreviations and Acronyms

AAV: Adeno-associated virus
ACSL: Acyl-CoA synthetase long-chain
AKT: AKT serine/threonine kinase
CD80: Cluster of differentiation 80
CD86: Cluster of differentiation 86
CD209: Cluster of differentiation 209
Ea: Effective arterial elastance
Ees/Ea: Ratio of End-systolic elastance/effective arterial elastance
CCL: C-C motif chemokine ligands
CXCL: C-X-C motif chemokine ligands
DAMP: Damage associated molecular patterns
DAPI: 6-diamidino-2-phenylindole
GFP: Green fluorescent protein
HIF: Hypoxia inducible factor
HMOX: Heme oxygenase 1
KEGG: Kyoto encyclopedia of genes and genomes
LPC: Lysophosphatidylcholines
LPE: Lysophosphatidylethanolamine
LPS: Lysophosphatidylserine
MCT: Monocrotaline
MerTK: Mer proto-oncogen, tyrosine kinase
mPAP: Mean pulmonary arterial pressure
MS: Mass spectrometry
NLRP3: NOD-, LRR-, and pyrin domain-containing protein 3
NODAL: Nodal growth differentiation factor
PAEC: Pulmonary arterial endothelial cell
PAH: Pulmonary arterial hypertension
PH: Pulmonary hypertension
PASMC: Pulmonary artery smooth muscle cells
PC: Phosphatidylcholines
PCBP2: Poly(rC)-binding protein 2
PE: Phosphatidylethanolamine
PI3K: Phosphoinositide 3-kinase
PS: Phosphatidylserine
PVR: Pulmonary vascular resistance
RNA-seq: RNA-sequencing
ROS: Reactive oxygen species
RV: Right ventricular/right ventricle
RVSP: Right ventricular systolic pressure
SNP: Single-nucleotide polymorphisms
TAPSE: Tricuspid annular plane systolic excursion
VDAC: Voltage-dependent anion-selective channel
VEH: Vehicle

## Acknowledgements

We thank the Center for Metabolomics and Proteomics and the Genomics Center at the University of Minnesota for providing services related to the generation of RNAseq and proteomics data.

## Disclosures

KWP received grant funding from Bayer unrelated to this manuscript.

## Sources of Funding

SZP is funded by an American Heart Association Career Development Award (23CDA1049093, https://doi.org/10.58275/AHA.23CDA1049093.pc.gr.167948) and by NIH K08 HL168166. ELB is funded by NIH R01s FD007627, HL146588, HL163960, HL155278, DK124845, and NIH R61 HL158941. This work was supported by the NIHR BioResource which supports the UK National Cohort of Idiopathic and Heritable PAH; the British Heart Foundation (BHF SP/12/12/29836) and the UK Medical Research Council (MR/K020919/1). CJR is supported by BHF Basic Science Research fellowship (FS/SBSRF/21/31025). JWW is funded by NIH R01s HL166843 and AI165553. KWP is funded by NIH R01s HL158795 and HL162927. The Orbitrap Eclipse instrumentation platform used in this work was purchased through High-end Instrumentation Grant S10OD028717 from the NIH.

---

**Central Figure.**
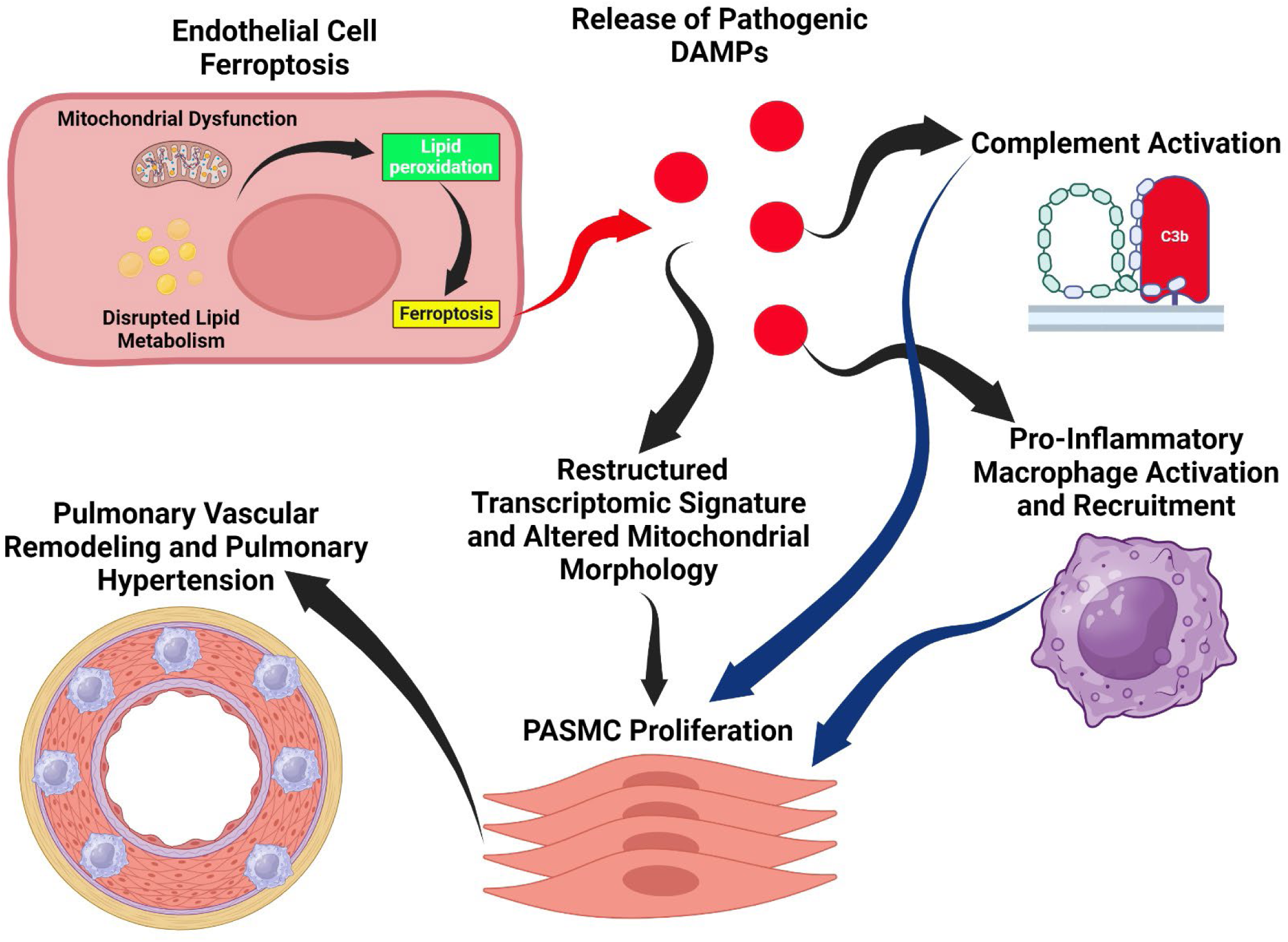
Proposed Model for the Pathogenic Role of Endothelial Cell Ferroptosis in PAH Pathobiology.

## Novelty and Significance

What is known?

- Ferroptosis is a mode of cell death initiated by lipid peroxidation which results from altered mitochondrial function. Ferroptotic death is highly inflammatory as it leads to release of DAMPs that activate the innate immune system
- PAH is a complex disease marked by mitochondrial derangements in endothelial and pulmonary artery smooth muscle cells, immune cell infiltration, and ectopic complement activation, but a mechanism that integrates all these pathogenic phenotypes is lacking.

What new information does this article contribute?

- Small molecule-mediated ferroptosis inhibition mitigates PAH severity even when evaluated in a translational approach, and multi-omic analyses demonstrate ferroptosis promotes PAH by altering the cellular architecture with increases in PASMC and interstitial macrophage relative abundances, restructuring the lung metabolic landscape, and enhancing peri-vascular complement deposition.
- Ferroptotic DAMPs from human PAEC alter human PASMC genetic regulation and impart changes that phenocopy many previously described PASMC phenotypes including disruptions in mitochondrial morphology, suppression of transcripts responsible for DNA repair and metabolism, activation of HIF/mTOR, and heightened proliferation rates.
- Ferroptotic DAMPs from human PAEC activate monocytes and promote a pro-inflammatory phenotype characterized by higher expression of CD86 and CD209 and lower expression of MerTK.
- AAV1-*Acsl4* treatment results in an inflammatory PAH phenotype in rodents with ectopic complement deposition, suggesting endothelial cell ferroptosis may be sufficient to induce pulmonary hypertension.
- SNPs in ferroptotic genes are independently associated with pulmonary hypertension severity in the Vanderbilt BioVU database.

## Notes

### Summary of Updates

We have updated the manuscript to include new data and improve figure quality.

