## Supplemental Data for "Ferroptosis Integrates Mitochondrial Derangements and Pathological Inflammation to Promote Pulmonary Hypertension"

**Supplemental Methods**

*Rodent Studies*: Small-molecule mediated ferroptosis inhibition in the monocrotaline (MCT rat PAH model was evaluated in male Sprague-Dawley rats. Rodents were randomly assigned to one of three groups: 1. Control rats injected with phosphate buffered saline (*n*=5), 2. Rats injected with 60 mg/kg MCT and then treated with daily intraperitoneal injections of vehicle (2% dimethyl sulfoxide, 50% polyethylene glycol, 5% Tween 80, and 43% double distilled water) starting two weeks after MCT injection (*n*=10), and 3. MCT rats treated with daily intraperitoneal ferrostatin-1 (1 mg/kg, Selleck Chemicals) starting two weeks after MCT injection (*n*=10). End-point analysis was performed 24 days post MCT injection. AAV1-GFP and AAV1-r*Acsl4* (produced by Vector Biolabs) driven by the cadherin-5 promoter at a dose of $0.3x10^{11}$ vector genomes were delivered via intratrachael injection the same day as low dose (30 mg/kg, Sigma-Aldrich) monocrotaline. The end-point studies for viral study were conducted 24 days post treatment. All rodent studies were approved by the University of Minnesota Institutional Animal Care and Use Committee

*Cardiovascular Phenotyping*: A Vevo2100 ultrasound system was used to perform echocardiographic analysis at the University of Minnesota Imaging Center[1-4]. Closed-chest pressure-volume loop analysis using a high-fidelity Millar 1.9F catheter (measured with Transonic Systems) was used to evaluate RV volumes and pressures. Pressure-volume loop measurements were analyzed on a LabScribe software system[1-4].

*Histological Examination of Pulmonary Vasculature*: Histological analysis examined percent medial thickness of *n*=4 control, *n*=4 MCT-Vehicle, *n*=4 MCT-Fer-1, *n*=5 AAV1-GFP, and *n*=5 AAV-*Acsl4* pulmonary vessel wall size. Lung tissue was fixed in 10% formalin, paraffin embedded, sectioned at 4-μm, and stained with hematoxylin and eosin at the University of Minnesota Histology and Research Laboratory. All images were collected by KWP or SZP and blindly analyzed by NTV and BK using FIJI (NIH).

*Histological Examination of RV Cardiomyocyte Cross Sectional Area*: Histological analysis examined RV cardiomyocyte cross sectional area of *n*=5 AAV1-GFP, and *n*=5 AAV-*Acsl4* pulmonary vessel wall size. RV free wall specimens were was in phosphate buffered saline and then immediately fixed in 10% formalin. We evaluated 2-4 areas of each heart and only cardiomyocytes that were circular were assessed when determining cross sectional area. Sections were stained with hematoxylin and eosin at the University of

Minnesota Histology and Research Laboratory as previously described[3]. All images were collected by SZP and blindly analyzed by BK using FIJI.

*Mitochondrial Proteomics Analysis*: Mitochondrial enrichments from lung tissue were subjected to TMT16-plex (ThermoFisher Scientific) labeling and quantitative proteomics using Proteome Discover Software as previously described[1,4,5]. We evaluated the lung mitochondrial-enriched proteome in *n*=5 control, *n*=5 MCT-Vehicle, and *n*=6 MCT-Fer-1 animals.

*Cell Culture*: Human PASMC (Lonza CC-2581) were grown with media and supplements (Lonza CC-3182) and passaged with subculture reagents (Lonza CC-5034). PASMC cells are shipped at passage 2 (P2) and were used at P4-6. Human PAEC (ATCC PCS-100-022) were grown in media (ATCC PCS1000030) with supplements (ATCC PCS100041) and passaged with reagents (ATCC PCS999003, PCS999004). PAEC cells arrived at P1 and were used at P3-6. Erastin 40 μM (ApexBio B1524) was added to nearly confluent PAEC cells for 3d. Ferrostatin-1 (Fer-1, Selleck Chemicals S7243), 4 μM, was added to spent media from erastin-treated and vehicle-treated PAEC. Spent media was applied to PASMC or macrophage cells in a 1:1 mix with PASMC/macrophage media for 24h.

*PASMC Mitochondrial Morphological Analysis*: After incubation with ferroptotic PAEC media or control PAEC media, PASMC were stained with MitoTracker Orange (Thermo Scientific, M7514), fixed in 4% paraformaldehyde, and then mounted in Prolong Mountant with NucBlue (Thermo Scientific). Images were collected on a Zeiss LSM 900 Airyscan 2.0 microscope and then blindly analyzed by RM using the Mitochondria Network Analysis Plug-In[6] (https://github.com/StuartLab) in FIJI.

*PASMC Replication Analysis*: PASMC incubated with ferroptotic PAEC media or control PAEC media for 24 hours and then were fixed in 4% PFA, solubilized with 1% Triton X-100 in PBS, washed/blocked with 5% goat serum and then incubated with a Ki67 antibody (Abcam, ab15580) at a 1:50 dilution overnight at $4^{0}$C. Cells were then washed/blocked with 5% goat serum in PBS, incubated with Alexa Fluor 568 anti-rabbit antibody (Invitrogen 1:500). Samples were washed with PBS, exposed to Hoechst stain, and mounted in Prolong Glass Antifade Mountant. Images were collected on a Zeiss LSM 900 Airyscan 2.0 microscope using identical settings and blinded analyzed by RTM to identify the proportion of Ki67 positive nuclei using FIJI. Cells were stained with secondary antibody alone to confirm immunoreactivity of Ki67 antibody.

*Monocyte Isolation and Analysis*: Monocytes from one donor were isolated from 20 mLs of whole blood using EasySep Direct Human Monocyte Isolation Kit (Stemcell Technologies, Cat# 19669) per the manufacturer's protocol. Following isolation, purified monocytes were seeded at 200,000 cells/well in a 12 well plate. Cells were then cultured in either 50% monocyte media (RPMI, 10% FBS, 1% Pen-strep) and 50% untreated PAEC supernatant or 50% monocyte media with 50% supernatant from erastin treated PAEC overnight. Fer-1, 2 μM, was added to eliminate primary effects of erastin. After overnight incubation, the cells were lifted from plates using scraping then counted and stained with flow cytometry antibodies at 1:100 dilution. Cells were analyzed using a 5 laser Cytek Aurora. The following antibodies were used: Anti-Human CD45 AF700 (BD Bioscience, Clone: HI30), Anti-Human CD64 APC-Cy7 (ThermoFisher, Clone: 10.1), Anti-Human CD86 BV605 (Biolegend, Clone:BU63), Anti-Human CD80 PE Cy-7 (Biolegend, Clone:2D10), Anti-Human CD200R PE (Biolegend, Clone:OX-108), Anti-Human MerTK BV711 (Biolegend, Clone:590H11G1E3).

*RNA sequencing of lung extracts and PASMC*: RNA from rat lung tissue was isolated from *n*=4 control, *n*=4 MCT-Vehicle and *n*=4 MCT-Fer-1 using PureLink RNA Mini kit (Thermo Fisher) with DNase. RNA from PASMC was collected using the same kit. Concentrations of RNA were determined through spectrophotometry by NanoDrop 2000. RNA sequencing and library preparations were conducted at the University of Minnesota Genomics Center via Illumina NovaSeq 6000 with 20 million reads per sample. Overall quality of RNAsequencing data was examined by hierarchical cluster analysis. One MCT-Fer-1 treated animal was a significant outlier (Supplemental Figure 6) and this sample was not used for correlational analyses.

*PASMC RNA expression analysis*: PASMC were plated into 10 cm dishes. When 90% confluent, cells were treated with spent media from erastin-treated and vehicle-treated PAEC media to which Fer-1, 4 μM, was added. Spent media was applied 1:1 with PASMC media to PASMC cells for 24h. Cells from 4 separate dishes of each erastin/vehicle treated cells were then lysed and RNA was isolated using Purelink RNA Mini Kit (Invitrogen) according to kit instructions. RNA was submitted to University of Minnesota Genomics Center for library creation (TruSeq Stranded mRNA, converted to Aviti compatible with Element Adept Rapid PCR + kit) and greater than $250x10^6$ reads were generated by sequencing. R-studio was used to delineate differentially expressed transcripts using a fold change greater than 1.5 and a false discovery rate <0.05.

*Deconvolution RNAsequencing Analysis*: Single cell reference data Reference data (expression matrix and metadata) was downloaded from http://mergeomics.research.idre.ucla.edu/PVDSingleCell/CellBrowser/?ds=PAHRatLungs#[7]. The expression matrix had the transformed ln(counts+1) values. Only the libraries corresponding to MCT and control treatments were used for further analysis. Single cell datasets for each treatment were checked for quality and cell composition. Marker genes for each cell type in both reference datasets were found using MAST and the Seurat FindMarkers functions. Since MAST provided a wider range of gene markers, this option was chosen. A reduced expression signature matrix for each celltype and condition was created using the DWLS R package (sigMatrix.MAST.r). Bulk RNAseq preparation: Fastq files were trimmed, filtered for low quality reads, aligned to the reference rat genome (Rattus_norvegicus-109) and summarized in a raw counts matrix (genes by samples) using the CHURP-PURR pipeline. Deconvolution**:** Bulk raw counts were used without any normalization or further filtering as suggested[8]. Control and MCT BulkRNA seq libraries were deconvolved individually using the signature matrix prepared with the corresponding reference dataset. Cell-type specific differential gene expression: Bulk raw counts were processed before differential gene expression testing. From the 30560 genes annotated in the reference genome, only those longer than 300 bp (25680) and with at least 10 cpm per treatment (control, MCT-Veh or MCT-Fer-1) were kept. This resulted in a dataset containing 15693 genes. The cell type proportions estimated with DWLS were combined to reduce the number of variables and allow the construction of the gene expression model for DE testing. The processed counts and combined cell type proportions were used to infer differentially expressed genes (DEGs) using the TOAST R package. Even though, individually some genes were detected as DE (*p*-value <0.05), when applying the BH multiple testing correction, no differentially expressed genes were found by the algorithm using a false discovery rate threshold of 0.05. Gene set enrichment analysis (GSEA): Despite not having enough power to reach significant results, the gene list ranked by descending FDR value was used for GSEA against the Hallmark gene sets from the MSigDb, which comprise 7300 genes organized in 50 gene sets. This gene list was trimmed so that genes not used in the DEG testing would not be included in the gene sets leaving a total of 6683. Set enrichment was tested using the ClusterProfiler package and the following options:

method=fgsea, exponent = 1, nPermSimple = 100000. Sets were considered enriched if most of their genes were located towards the end of the ranked gene list (i.e. low FDR values from the DE test); therefore, an additional filtering step was performed on the GSEA results by only considering sets with negative NES<0 and FDR<0.05. Additionally, GSEA results of enriched pathways were individually inspected using their enrichment plots.

*Lung Metabolomics Analysis*: Global metabolomics analysis of frozen lung specimens on *n*=5 Control, *n*=5 MCT-Vehicle, and *n*=5 MCT-Fer-1 was performed at the University of Minnesota Center for Metabolomics and Proteomics using the Biocrates' MxP® Quant 500 kit. 100 mg of sample was placed in 2.0 mL Precelly standard tubes and homogenized 3 times for 30 seconds at 5,800 rpm. Samples were centrifuged at 10,000*g* for 5 minutes at 4°C and supernatant was collected. 10µL of the extract was loaded onto a well insert. Classes of metabolites were determined using 50µL of 1:1:1:0.16 water:EtOH:pyridine:phenyl isothiocyanate solution and incubated for an hour. ABSciex QTRAP 5500 triple-quadrupole (Farmington, MA, USA) mass spectrometer was used to perform metabolomic assays.

*Confocal Microscopy Analysis of Lung Sections*: Confocal microscopy was performed on *n*=4 Control, *n*=4 MCT-Vehicle and *n*=4 MCT Fer-1 or *n*=5 AAV1-GFP and *n*=5 AAV1-*Acsl4* treated animal lung sections to evaluated perivascular complement deposition, CD11b cellular reactivity (ferrostatin treatment arm), and ACSL4 (AAV1-GFP and AAV1-*Acsl4* rats) immunoreactivity. Lung tissue slides were de-paraffinized through xylene and ethanol washes, heated in Reveal Decloaker Buffer (BioCare Medical, Pacheco, CA) for 30 minutes. Sections were washed/blocked with 5% goat serum in PBS and then and stained with primary antibodies to C3 (Abcam, ab200999) or CD11b (Thermo Scientific, PA5-90724) and FITC conjugated alpha-smooth muscle actin (Sigma Aldrich, 1A4) at a 1:50 dilution in goat serum. Slides were washed then incubated anti-rabbit AlexaFluor-568 antibody (Thermo Fisher) at a dilution of 1:500 for 30 minutes at 37°C. Samples were washed with PBS, exposed to Hoechst stain, and treated with an autofluorescence quenching kit (Vector Laboratories). Finally, samples were mounted in Prolong Glass Antifade Mountant (Thermo Scientific). Images were collected on a Zeiss LSM 900 Airyscan 2.0 microscope using identical settings by KWP or RM and blindly analyzed by NTV or RM. Relative fluorescence intensity normalized to area was calculated in FIJI (NIH) and compared across the three experimental groups. For CD11b analysis, we quantified the number of CD11b positive nuclei per arteriole. For each antibody, sections were stained with secondary antibody alone to confirm immunoreactivity of primary antibodies.

*Human Genetics Analysis*: Vanderbilt University de-identified electronic health record database and associated DNA biobank, BioVU, was used to determine how the presence of single nucleotide polymorphisms (SNP) in ferroptosis genes were associated with PH severity. We searched the ClinVar database to identify SNPs in ferroptosis genes there were classified as either pathogenic or of uncertain significance. SNPs were then searched in BioVU and the impact of the presence of SNPs on echocardiography and hemodynamic measures of pulmonary hypertension were evaluated (Supplemental Table 2). Human studies were approved by the Vanderbilt University Institutional Review Board. The small number of SNP carriers were pooled ($n$=40 with 1 patient harboring 2 SNPs) for multivariate analysis and then multivariate linear regression analysis was performed to determine how the presence of SNPs associated with RVSP, mPAP, and PVR after adjusting for age, sex, race, use of PAH-specific therapy, the presence of chronic obstructive lung disease and heart failure.

*Statistics*: Statistical analyses were performed using Prism 9.0 (GraphPad Software). Normality of data was determined using the Shapiro-Wilk test. If data were normally distributed and there was equal variance as determined using the Brown-Forsythe test, 1-way analysis of variance with Tukey's multiple-comparisons test was performed. If there was unequal variance, Brown-Forsythe and Welch analysis of variance with Dunnett multiple-comparisons test was completed. If the data were not normally distributed, the Kruskal-Wallis test and Dunn's multiple-comparisons test were used when comparing two groups and the Mann Whitney U test for comparing two groups. Graphs display the mean or median value and all individual values. $P$-values of <0.05 were considered to indicate statistical significance. Hierarchical cluster analyses, sparse least discriminate analysis, random forest classification, and correlational heatmapping were performed using MetaboAnalyst software. The relative abundance of each protein was determined using Proteome Discover Software. Transcripts and proteins that were significantly correlated ($p$<0.05) with right ventricular systolic pressure and end-arterial elastance (Ea) were processed for Kyoto Encyclopedia of Genes and Genomes (KEGG) using ShinyGO 0.76.3 (http://bioinformatics.sdstate.edu/go/).

Supplemental Figure 1: Transcriptomic correlates of RVSP and associations between ferroptosis transcripts and RVSP.

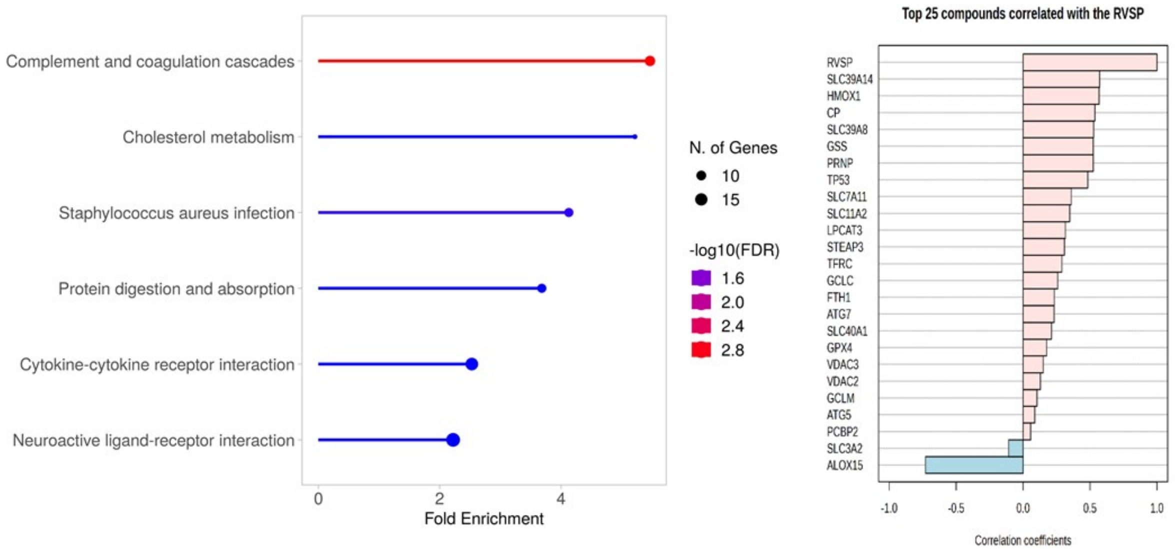

Supplemental Figure 2: Combined Metabolomics/Lipidomics Profiling of Whole Lungs Identified Changes in Lipid Metabolism with Ferrostatin-1**.** (A) Hierarchical cluster analysis of the most divergently regulated 250 metabolites in the lungs. (B) Sparse least squares discriminant analysis highlighted differences in the three experimental groups. (C) Random forest classification identified the top 15 most important metabolites for differentiating the three groups. Lipid species dominated the changes in the lung. (D) Hierarchical cluster analysis of lysophophatidylcholines (LPC) and phosphitidylcholines (PC). Ferrostatin-1 treatment increased a subpopulation of LPC and PC. (E) Hierarchical cluster analysis showed ferrostatin-1 reduced almost all LPE/PE species. Lung LPS+PS (F) and amino acids (G) levels were minimally altered by ferrostatin-1 treatment as compared to MCT-Vehicle. CE: Cholesterol ester, HxCer: Hexosylceramide, PE: phosphatidylethanolamine, LPC: Lysophophatidylcholine, PC: Phosphatidylcholine, PS: Phosphatidylserine, PA: Phosphatidic acid.

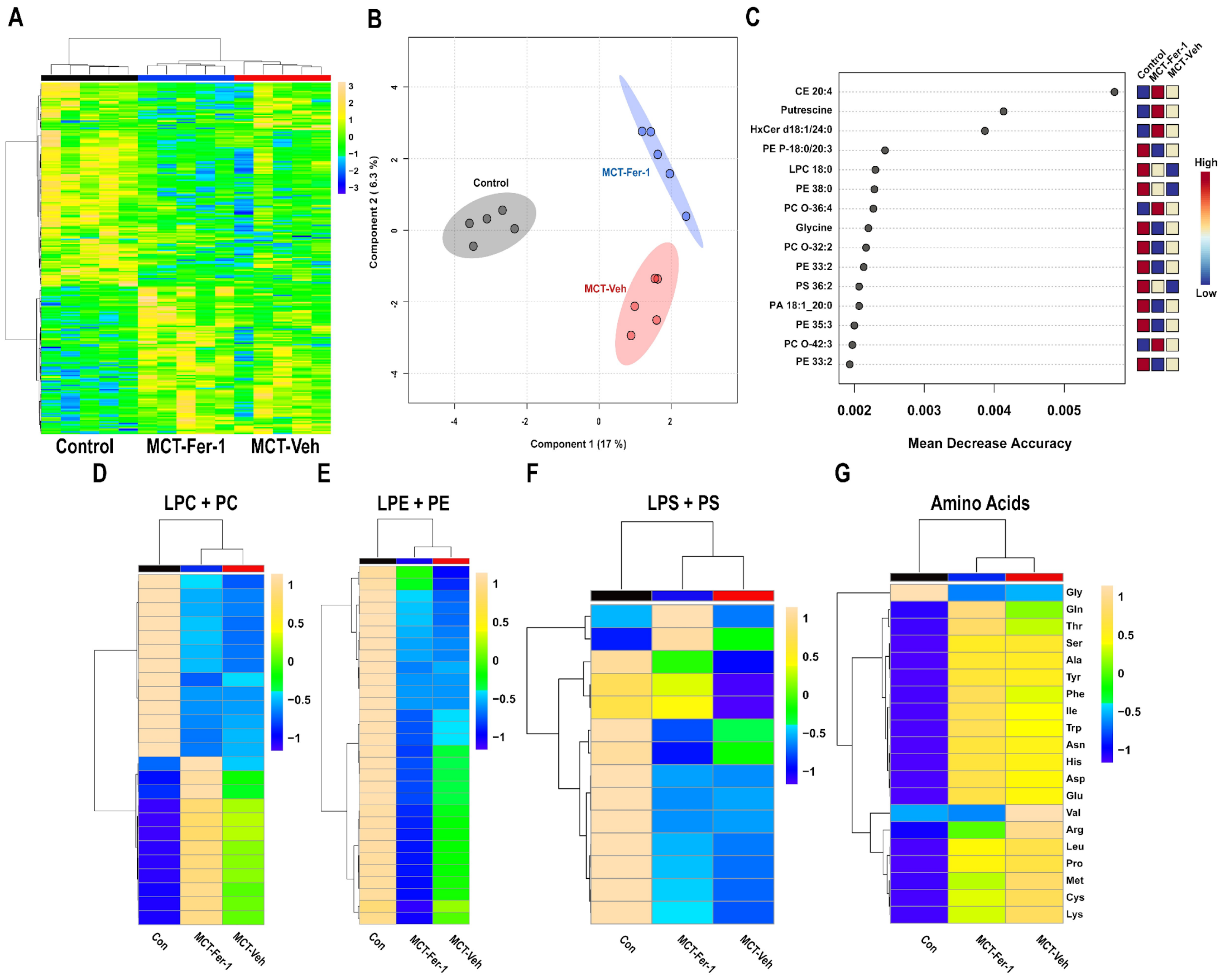

A
Control
MCT-Fer-1
MCT-Veh
B
Control
MCT-Fer-1
MCT-Veh
Component 1 (17 %)
Component 2 ( 6.3 %)
C
Control
MCT-Fer-1
MCT-Veh
CE 20:4
Putrescine
HxCer d18:1/24:0
PE P-18:0/20:3
LPC 18:0
PE 38:0
PC O-36:4
Glycine
PC O-32:2
PE 33:2
PS 36:2
PA 18:1_20:0
PE 35:3
PC O-42:3
PE 33:2
High
Low
Mean Decrease Accuracy
D
LPC + PC
E
LPE + PE
F
LPS + PS
G
Amino Acids
Gly
Gln
Thr
Ser
Ala
Tyr
Phe
Ile
Trp
Asn
His
Asp
Glu
Val
Arg
Leu
Pro
Met
Cys
Lys
Con
MCT-Fer-1
MCT-Veh

Supplemental Figure 3: Ferrostatin-1 treatment modulates the cytokine/chemokine profile of the lungs. (A) Hierarchical cluster analysis of cytokine/chemokine pathway from RNA-seq counts. (B) Random forest classification-based identification of transcripts in the cytokine/chemokine pathway that differentiated the three experimental groups. (C) 15 transcripts that most strongly associated with Ea and their respective *p*-values. (D) 15 transcripts that most strongly associated with RVSP and their respective *p*-values

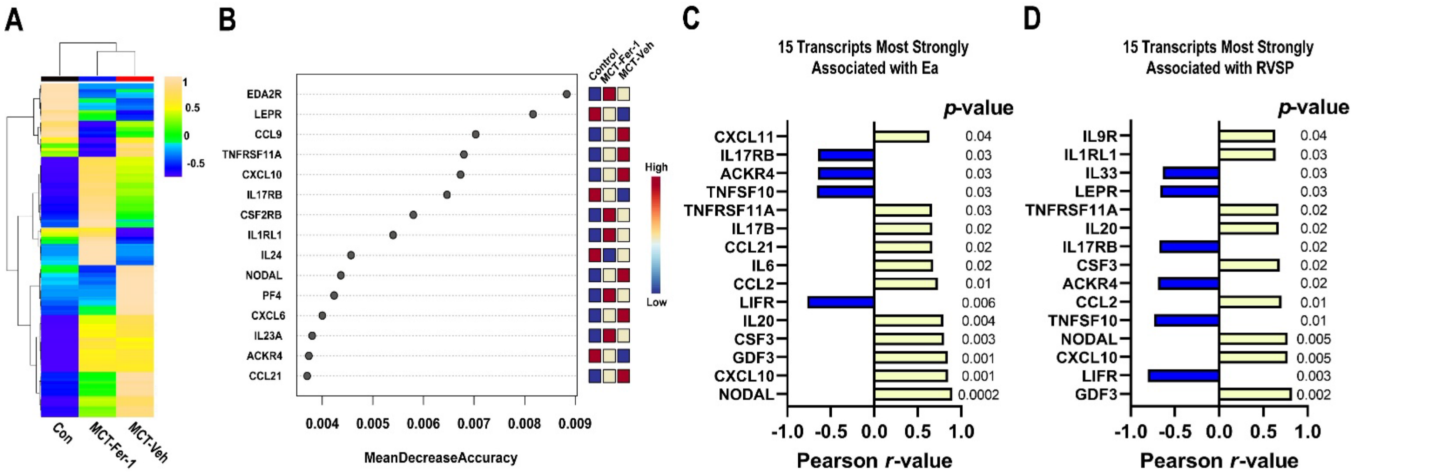

Supplemental Figure 4: Pathway Analysis of Upregulated Transcripts from Pulmonary Artery Endothelial Cells Analyzed by Single Cell RNA-seq. (A) KEGG pathway analysis of top 40 pathways identified from upregulated transcripts. (B) Wiki Pathway Analysis of upregulated transcripts in PAH endothelial cells.

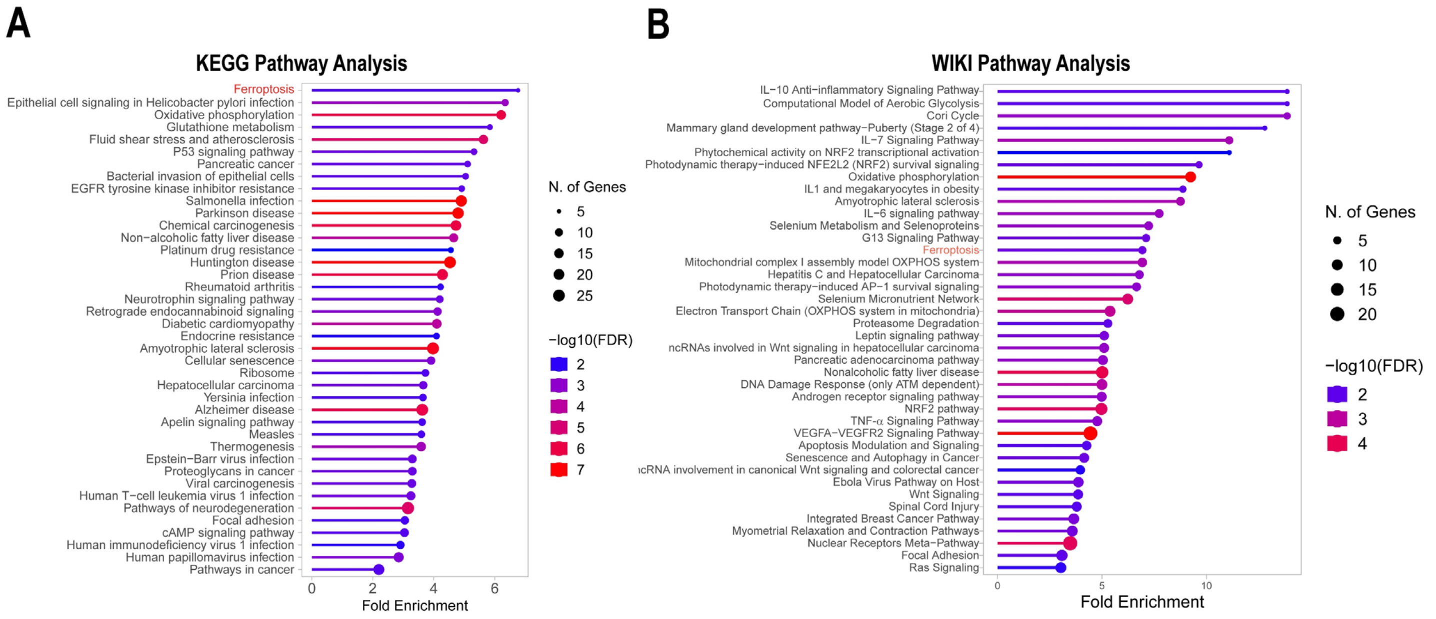

Supplemental Figure 5: Identified Ferroptotic SNPs Were Not Associated With PAH Severity in the NIHRBR dataset. Presence of ferroptotic SNPs were not associated with differences in (A) mPAP (Carrier: 38 mm Hg (33, 52), Noncarrier: 50 mm Hg (41, 60), *p*=0.13) or (B) PVR (Carrier: 10.9 Wood units (4.4, 16.3), Noncarrier: 10.9 Wood units (7.7, 15.1), *p*=0.68). *p*-values determined by Mann-Whitney test.

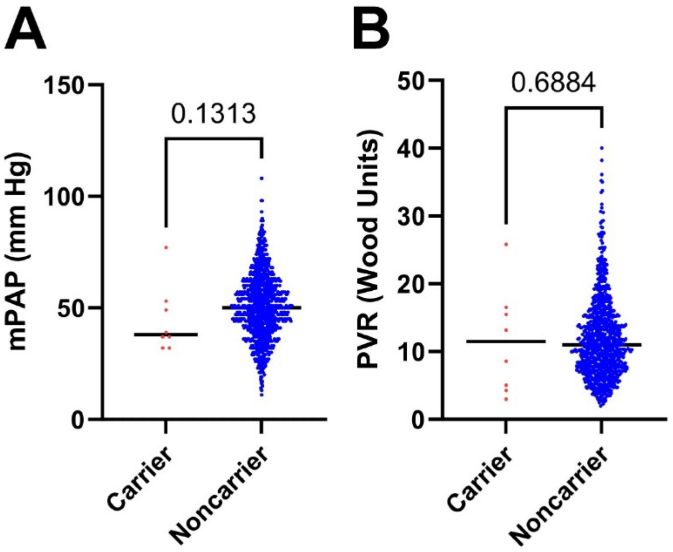

Supplemental Figure 6: Hierarchical cluster analysis of RNA-sequencing data shows ferrostatin-1 treated sample was a significant outlier.

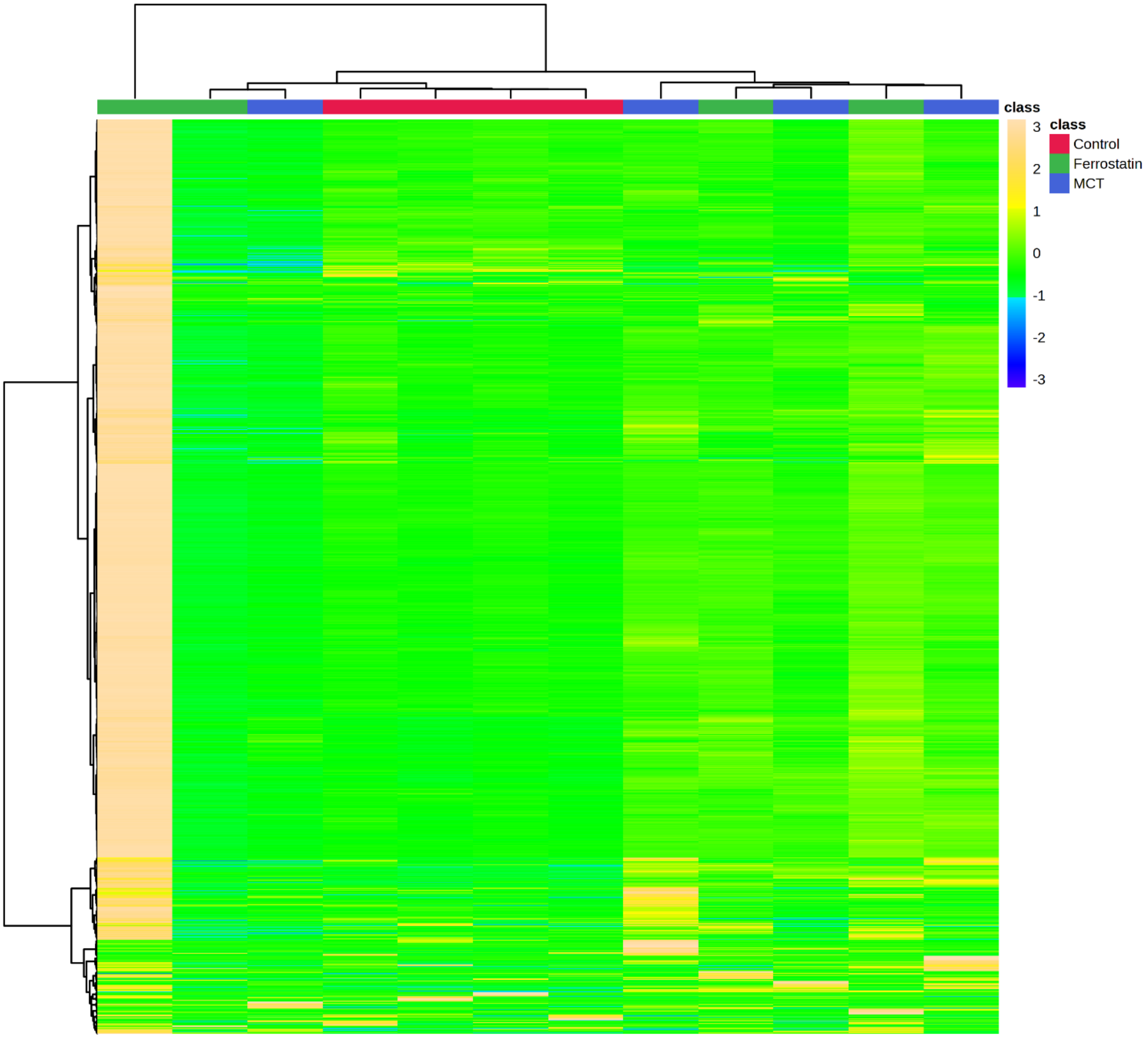

Supplemental Figure 7: Ferrostatin-1 treatment slightly increased right ventricular systolic pressure in control animals but not to pathological levels.

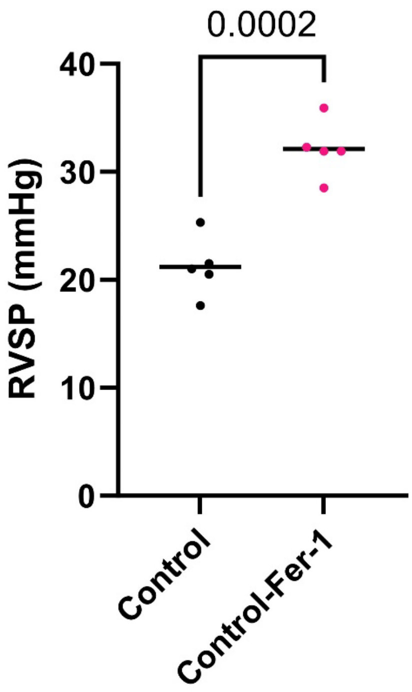

Supplemental Table 1: Summary of Cell Specific Changes Identified in Deconvolution RNAseq Experiments

| cell_type | avg_control | sd_control | avg_MCT | sd_MCT | avg_ferrostatin | sd_ferrostatin |
|---|---|---|---|---|---|---|
| AT2_cells | 22.929 | 0.758 | 18.503 | 3.709 | 18.244 | 3.118 |
| Endothelial_arterial_1 | 18.441 | 2.100 | 10.552 | 2.853 | 13.318 | 1.590 |
| AT1_cells | 15.883 | 1.735 | 11.441 | 2.113 | 12.457 | 0.870 |
| Smooth_muscle_cells | 7.367 | 0.867 | 9.745 | 0.746 | 8.720 | 0.403 |
| Club_cells | 7.177 | 0.633 | 5.647 | 2.920 | 5.147 | 1.219 |
| Endothelial_capillary | 6.988 | 0.561 | 6.990 | 1.015 | 8.425 | 0.490 |
| Mesothelial_cells | 3.285 | 0.521 | 3.145 | 1.174 | 3.142 | 0.386 |
| Endothelial_arterial_2 | 3.085 | 0.955 | 5.662 | 1.182 | 5.134 | 1.360 |
| Fibroblasts | 3.034 | 0.687 | 4.302 | 0.698 | 3.966 | 0.631 |
| Alveolar_macrophages | 2.310 | 0.707 | 5.899 | 3.800 | 7.651 | 1.941 |
| B_cells | 1.765 | 0.536 | 0.979 | 1.162 | 0.415 | 0.641 |
| Interstitial_macrophages | 1.704 | 0.426 | 6.098 | 1.712 | 4.129 | 0.960 |
| Ciliated_cells | 1.093 | 0.161 | 1.885 | 1.348 | 1.478 | 0.532 |
| Plasmacytoid_dendritic | 0.863 | 0.087 | 0.967 | 1.050 | 1.018 | 0.534 |
| Conventional_dendritic | 0.808 | 0.540 | 1.628 | 1.897 | 0.745 | 1.490 |
| Non_classical_monocytes | 0.790 | 0.378 | 0.636 | 0.586 | 1.767 | 1.346 |
| Neutrophils | 0.660 | 0.473 | 0.690 | 0.830 | 0.382 | 0.763 |
| Classical_monocytes | 0.452 | 0.177 | 0.634 | 0.733 | 0.335 | 0.670 |
| Proliferating_macrophages | 0.389 | 0.188 | 1.385 | 1.607 | 1.410 | 1.090 |
| NK_cells_1 | 0.312 | 0.257 | 0.172 | 0.288 | 0.096 | 0.117 |
| Mast_cells | 0.228 | 0.183 | 1.469 | 0.632 | 1.363 | 0.315 |
| T_cells_Serpinb6 | 0.211 | 0.275 | 0.463 | 0.535 | 0.136 | 0.273 |
| ILC2 | 0.134 | 0.090 | 0.088 | 0.132 | 0.072 | 0.145 |
| CD8_T_cells | 0.092 | 0.184 | 0.000 | 0.000 | 0.199 | 0.398 |
| Naive_T_cells | 0.000 | 0.000 | 0.500 | 0.578 | 0.053 | 0.106 |
| NK_cells_2 | 0.000 | 0.000 | 0.000 | 0.000 | 0.000 | 0.000 |
| Proliferating_T_cells | 0.000 | 0.000 | 0.084 | 0.110 | 0.124 | 0.249 |
| Regulatory_T_cells | 0.000 | 0.000 | 0.435 | 0.546 | 0.073 | 0.146 |

Supplemental Table 2: Relationship Between Individual Ferroptosis SNPs and Pulmonary Hypertension Severity in BioVU Database

**SNP:** rs78298849 **Gene:** *SLC3A2* **Effect:** Missense: I163V

| | N | Carrier<br>*N=8* | Noncarrier<br>*N=2439* | *p*-value |
|---|---|---|---|---|
| TRV | 2444 | 3.0 **3.6** 3.7<br>3.6±0.6 | 2.7 **3.1** 3.7<br>3.2±0.8 | *p*=0.13[1] |
| RVSP | 2344 | 43.5 **60.4** 68.0<br>59.7±18.2 | 35.0 **47.0** 62.7<br>51.7±23.3 | *p*=0.151[1] |
| mPAP | 2282 | 27.2 **38.0** 50.2<br>39.1±16.0 | 22.0 **31.0** 41.0<br>32.5±14.0 | *p*=0.209[1] |
| PVR | 2447 | 2.9 **4.5** 5.5<br>6.4±7.4 | 1.7 **2.6** 4.6<br>4.0±4.1 | *p*=0.157[1] |

**SNP X:108902750** **Gene:** ***ACSL4*** **Effect:** Intron Variant

| | N | Carrier (Two Copies)<br>*N=2* | Carrier (One Copy)<br>*N=11* | Noncarrier<br>*N=2429* | *p*-value |
|---|---|---|---|---|---|
| TRV | 2444 | 3.6 **3.8** 3.9<br>3.8±0.6 | 3.2 **3.7** 4.3<br>3.7±0.8 | 2.7 **3.1** 3.7<br>3.2±0.8 | *p*=0.013[1] |

| | | | | | |
|---|---|---|---|---|---|
| RVSP | 2344 | 64.5 **69.0** 73.5<br>69.0±12.7 | 45.0 **55.0** 79.5<br>63.2±24.0 | 35.0 **47.0** 62.7<br>51.7±23.3 | *p*=0.03[1] |
| mPAP | 2282 | 44.3 **46.5** 48.8<br>46.5±6.4 | 21.0 **28.0** 40.5<br>30.8±14.0 | 22.0 **31.0** 41.0<br>32.5±14.0 | *p*=0.772[1] |
| PVR | 2447 | 6.5 **7.3** 8.2<br>7.3±2.5 | 2.4 **3.2** 5.4<br>4.4±3.7 | 1.7 **2.6** 4.6<br>4.0±4.1 | *p*=0.195[1] |

**SNP: rs8177318** **Gene:** ***TF*** **Effect:** Missense: S55R

| | **N** | **Carrier**<br>***N=3*** | **Noncarrier**<br>***N=2444*** | ***p*-value** |
|---|---|---|---|---|
| TRV | 2444 | 3.2 **4.2** 5.0<br>4.1±1.9 | 2.7 **3.1** 3.7<br>3.2±0.8 | *p*=0.38[1] |
| RVSP | 2344 | 54.8 **85.0** 112.5<br>83.2±57.8 | 35.0 **47.0** 62.7<br>51.7±23.2 | *p*=0.345[1] |
| mPAP | 2282 | 38.0 **45.0** 49.0<br>43.0±11.1 | 22.0 **31.0** 41.0<br>32.5±14.0 | *p*=0.141[1] |
| PVR | 2447 | 3.7 **5.5** 5.5<br>4.3±2.1 | 1.7 **2.6** 4.6<br>4.0±4.1 | *p*=0.391[1] |

**SNP: rs139633388** **Gene:** ***CP*** **Effect:** Missense: G895A

| | N | Carrier<br>*N=4* | Noncarrier<br>*N=2435* | *p*-value |
|---|---|---|---|---|
| TRV | 2444 | 2.8 **3.7** 4.6<br>3.7±1.2 | 2.7 **3.1** 3.7<br>3.2±0.8 | *p*=0.463[1] |
| RVSP | 2344 | 44.2 **65.2** 88.6<br>67.6±35.4 | 35.0 **47.0** 62.7<br>51.7±23.3 | *p*=0.34[1] |
| mPAP | 2282 | 34.5 **46.0** 52.5<br>42.7±18.2 | 22.0 **31.0** 41.0<br>32.5±14.0 | *p*=0.259[1] |
| PVR | 2447 | 3.2 **6.0** 9.6<br>6.6±4.1 | 1.7 **2.6** 4.6<br>4.0±4.1 | *p*=0.081[1] |

**SNP: 3:148901395:TA** **Gene:** ***CP*** **Effect:** Intron Variant

| | N | Carrier<br>*N=9* | Noncarrier<br>*N=2362* | *p*-value |
|---|---|---|---|---|
| TRV | 2444 | 3.1 **3.6** 4.3<br>3.6±0.9 | 2.7 **3.1** 3.7<br>3.2±0.8 | *p*=0.124[1] |
| RVSP | 2344 | 50.0 **63.0** 75.2<br>66.6±28.7 | 35.0 **47.0** 62.7<br>51.7±23.3 | *p*=0.066[1] |

| | | | | |
|---|---|---|---|---|
| mPAP | 2282 | 30.0 **41.0** 50.0<br>40.4±19.5 | 22.0 **31.0** 41.0<br>32.5±14.0 | *p*=0.126[1] |
| PVR | 2447 | 3.1 **6.7** 8.5<br>7.9±7.4 | 1.7 **2.6** 4.6<br>4.0±4.1 | *p*=0.033[1] |

**SNP: rs150303869** **Gene:** ***CP*** **Effect:** Missense: N263S

| | **N** | **Carrier**<br>***N=4*** | **Noncarrier**<br>***N=2443*** | ***p*-value** |
|---|---|---|---|---|
| TRV | 2444 | 3.5 **3.9** 4.2<br>3.9±0.5 | 2.7 **3.1** 3.7<br>3.2±0.8 | *p*=0.05[1] |
| RVSP | 2344 | 63.5 **71.7** 78.8<br>70.5±14.9 | 35.0 **47.0** 62.7<br>51.7±23.3 | *p*=0.042[1] |
| mPAP | 2282 | 39.8 **49.5** 52.2<br>42.5±16.5 | 22.0 **31.0** 41.0<br>32.5±14.0 | *p*=0.166[1] |
| PVR | 2447 | 2.6 **4.8** 7.9<br>5.7±4.8 | 1.7 **2.6** 4.6<br>4.0±4.1 | *p*=0.389[1] |
| *a* ***b*** *c* represent the lower quartile *a*, the median *b*, and the upper quartile *c* for continuous variables.<br>*x*±*s* represents X±1 SD. *N* is the number of non-missing values. Tests used: [1]Wilcoxon test; [2]Pearson test. | | | | |

Abbreviations: TRV: Tricuspid regurgitation velocity, RVSP: right ventricular systolic pressure, PVR: pulmonary vascular resistance, MPAP: mean pulmonary arterial pressure

Supplemental Table 3: Characterization of BioVU patients harboring or lacking ferroptosis SNPs

| | ***N*** | **Noncarrier**<br>***N=2407*** | ***Carrier***<br>***N=40*** | ***p*-value** |
|---|---|---|---|---|
| Age | 2447 | 51.6 **61.3** 70.1<br>60.0±14.1 | 51.4 **57.0** 67.3<br>58.2±12.7 | *p*=0.265[1] |
| Sex | 2447 | | | *p*=0.099[2] |
| Female | | 47% | 60% | |
| Male | | 52% | 40% | |
| Race | 2447 | | | *p*=0.043[2] |
| Black | | 14% | 28% | |
| Other | | 2% | 3% | |
| White | | 84% | 70% | |
| BMI (kg/$m^2$) | 2387 | 24.8 **29.0** 34.2<br>30.8±17.5 | 24.0 **26.7** 33.8<br>29.1±6.4 | *p*=0.402[1] |
| RVSP (mm Hg) | 2344 | 35 **47** 62<br>52±23 | 47 **64** 79<br>65±26 | *p*<0.001[1] |

| | | | | |
|---|---|---|---|---|
| PVR (Wood units) | 2447 | 1.7 **2.6** 4.6<br>4.0±4.1 | 2.4 **4.3** 6.8<br>5.6±4.6 | *p*=0.001[1] |
| mPAP (mm Hg) | 2282 | 22 **31** 41<br>32±14 | 25 **40** 51<br>38±15 | *p*=0.017[1] |
| Hypertension | 2447 | 80% | 70% | *p*=0.112[2] |
| COPD | 2447 | 22% | 20% | *p*=0.778[2] |
| ILD | 2447 | 31% | 40% | *p*=0.241[2] |
| Heart Failure | 2447 | 71% | 68% | *p*=0.5872 |
| OSA | 2447 | 23% | 22% | *p*=0.9632 |

*a* ***b*** *c* represent the lower quartile *a*, the median *b*, and the upper quartile *c* for continuous variables.

*x* ± *s* represents X ± 1 SD. *N* is the number of non-missing values. Tests used: [1]Wilcoxon test; [2]Pearson test

Abbreviations: BMI: body mass index, RVSP: right ventricular systolic pressure, PVR: pulmonary vascular resistance, MPAP: mean pulmonary arterial pressure, COPD: Chronic obstructive lung disease, ILD: Interstitial Lung Disease, OSA: Obstructive sleep apnea
